# Verbalizing phylogenomic conflict: Representation of node congruence across competing reconstructions of the neoavian explosion

**DOI:** 10.1101/233973

**Authors:** Nico M. Franz, Lukas J. Musher, Joseph W. Brown, Shizhuo Yu, Bertram Ludäscher

## Abstract

Phylogenomic research is accelerating the publication of landmark studies that aim to resolve deep divergences of major organismal groups. Meanwhile, systems for identifying and integrating the novel products of phylogenomic inference – such as newly supported clade concepts – have not kept pace. However, the ability to *verbalize* both node concept congruence and conflict across multiple, (in effect) simultaneously endorsed phylogenomic hypotheses, is a critical prerequisite for building synthetic data environments for biological systematics, thereby also benefitting other domains impacted by these (conflicting) inferences. Here we develop a novel solution to the conflict verbalization challenge, based on a logic representation and reasoning approach that utilizes the language of Region Connection Calculus (RCC–5) to produce consistent *alignments* of node concepts endorsed by incongruent phylogenomic studies. The approach employs clade concept labels to individuate concepts used by each source, even if these carry identical names. Indirect RCC–5 modeling of *intensional* (property-based) node concept definitions, facilitated by the local relaxation of coverage constraints, allows parent concepts to attain congruence in spite of their differentially sampled children. To demonstrate the feasibility of this approach, we align two recently published phylogenomic reconstructions of higher-level avian groups that entail strong conflict in the “neoavian explosion” region. According to our representations, this conflict is constituted by 26 instances of input “whole concept” overlap. These instances are further resolvable in the output labeling schemes and visualizations as “split concepts”, thereby providing the provenance services needed to build truly synthetic phylogenomic data environments. Because the RCC–5 alignments fundamentally reflect the trained, logic-enabled judgments of systematic experts, future designs for such environments need to promote a culture where experts routinely assess the intensionalities of node concepts published by our peers – even and especially when we are not in agreement with each other.

## Introduction

Three years ago, Jarvis et al. (2014; henceforth 2014.JEA) [1] published a landmark phylogenomic reconstruction of modern, higher-level bird relationships. Within 12 months, however, another major yet differentially sampled analysis by Prum et al. (2015; henceforth 2015.PEA) [2] failed to support several of the deep divergences recovered in the preceding study, particularly within the Neoaves sec. (*secundum* = according to) Sibley et al. (1988) [3]. Thomas (2015) [4] used the term “neoavian explosion” to characterize the lack of congruence regarding these early-diverging bird tree inferences (see also [5]). Similarly, after reviewing six phylogenomic studies with reciprocally incongruent relationships, Suh [6] concluded that the root region of the Neoaves constitutes a “hard polytomy”. Multiple analyses have dissected the impact of differential biases in terminal and genome sampling, as well as evolutionary modeling and analysis constraints, on resolving this complex radiation [7, 8, 9]. Suh [6] argues that a well resolved consensus is not imminent (though see [10]). Brown et al. (2017) [11] analyzed nearly 300 avian phylogenies, but found somewhat unexpectedly that the most recent studies “continue to contribute new edges” to a group whose state of phylogenomic exploration is considered rather mature.

These recent advancements provide an opportunity to reflect on how synthesis should be realized in the age of phylogenomics [11, 12, 13]. The neoavian explosion can be considered a use case where multiple, almost simultaneously published studies provide strong, phylogenomically supported signals for conflicting hierarchies. Resolution towards a single, comprehensive, and universally adopted phylogeny is unlikely in the short term.

Rather than focusing on the analytical challenges along the path towards unitary resolution [9], we turn to the issue of how phylogenomic advancement with persistent conflict affects the technical and social design of synthetic, collaborative big data research infrastructures. Particularly *verbal* representations of the neoavian explosion are not well designed for big data integration in the face of persistent conflict [14]. Some authors use tree alignment graphs in combination with color and width variations to identify regions (edges) of phylogenomic congruence and conflict [15]. Other representations may show multiple incongruent trees side-by-side, while using consistent color schemes for congruent clade sections [9]. Yet others may use tanglegrams that are enhanced to highlight congruence [4], rooted galled networks [16] or neighbor-net visualizations [17] that show split networks for conflicting topology regions, or simply provide a consensus tree in which incongruent bifurcating branch inferences are collapsed into polytomy [6].

We hold that verbalizing phylogenomic congruence and conflict in use cases such as the neoavian explosion constitutes a novel representation challenge in systematics for which traditional solutions are inadequate. The aforementioned studies implicitly support this claim. All use largely overlapping sets of Code-compliant [18] and other higher-level names in the Linnaean tradition, with sources including [19] or [20]. To identify these study-specific name usages in the following discussion, we will utilize the *taxonomic concept label* convention of [14]. Accordingly, name usages sec. 2014.JEA are prefixed with “2014.”, whereas name usages sec. 2015.PEA are prefixed with “2015.”

We diagnose the verbalization challenge as follows. In some instances, identical clade names are polysemic – i.e., have multiple meanings – across studies. For instance, 2015.Pelecaniformes excludes 2015.Phalacrocoracidae, yet 2014.Pelecaniformes includes 2014.Phalacrocoracidae; reflecting in our representations on two incongruent meanings of “Pelecaniformes”. In other cases, two or more non-identical names are semantically congruent, e.g., 2015.Strisores and Caprimulgimorphae. Where author teams use names that are unique to just one study – e.g., Aequorlitornithes or 2014.Cursorimorphae – the meanings of these name usages are not always reconcilable without additional human effort, thereby adding an element of referential uncertainty to the apparent conflict. Lastly, many of the newly inferred edges are not named at all in representations of phylogenomic conflict [4]. There is an implicit bias towards labeling edges when suitable names are already available. Unnamed edges can create situations where conflict cannot be discussed or reconciled due to the lack of syntactic structure.

Jointly, the effects of polysemic names, synonymous names, exclusive yet hard-to-reconcile names, and conflicting unnamed edges are symptomatic of a data science culture that appears unprepared for the verbal representation challenges that phylogenomic studies present. Suppose we inquire how the author teams would build a collaborative, phylogenomically structured data environment that can individually represent and at the same integrate their conflicting hierarchies, from the tips to the root, and thus reliably respond to name-based data queries across these hierarchies. The naming system that each team uses individually is not suited for this task. Indeed, traditional naming approaches in systematics are context-constrained, and therefore under-designed for collaborative big data environments that can represent rapid advancement as well as persistent conflict in phylogenomic knowledge [21]. At root, this is a novel *conceptual* challenge for systematics, made imperative by the combination of accelerated generation of phylogenomic trees and creation of synthetic data environments that can ingest these for further integration and use [11, 13, 22, 23, 24]. The representation services that such environments aspire to provide require an appropriate implementation of *node identity* and *provenance*, and hence a conception of multi-node congruence or incongruence across individual trees and synthesis versions.

Here we propose a solution to the phylogenomic conflict representation challenge. The solution is an extension of prior concept taxonomy research [14, 25, 26], and deploys logic reasoning to align tree hierarchies based on Region Connection Calculus (RCC–5) assertions of node congruence [27, 28, 29]. We demonstrate the feasibility of this approach by aligning subregions and entire phylogenomic trees of the 2015./2014.Neornithes as inferred by 2015.PEA and 2014. JEA. In doing so, we address several broader representation challenges; such as the phylogenomically inferred paraphyly of classification schemes that are nevertheless said to be followed when labeling tree regions, or the inference of higher-level node congruence in spite of differentially sampled terminals. Based on the verbal and visual alignment products for the neoavian explosion use case, we answer the question of “how to build a synthetic data environment in the face of persistent phylogenomic conflict?” The subsequent discussion focuses on the feasibility and desirability of establishing big data synthesis services for the phylogenomic domain, with particular emphasis on the need to embrace trained expert judgment [30].

## Methods

### Syntactic and semantic conventions

Representing phylogenomic congruence and conflict makes it necessary to specify notions of identity and non-identity with regards to the terms, concepts, and relationships we use. Because our usages may differ somewhat from prevailing conventions, we will clarify these upfront.

We refrain in most instances from using the term “taxon” or “taxa”. We take taxa to constitute evolutionary entities whose members are manifested in the natural realm. The task for systematics is to successively approximate – via empirical inferential processes – the causally sustained identities and limits of these entities. Thus, we assign the status of ‘models’ to taxa, which systematists aim to ‘mimic’ with increasing precision and reliability in the realm of human empirical theory making. This perspective allows for realism about taxa, and also for the obvious need to let human-made representations *stand for* taxa [31], at any given time and however imperfectly, when the representations are needed to support inferences about evolutionary phenomena.

In reserving an external model status for taxa, we can create a pragmatically separate design space for the human language/theory making realm. To represent congruence and conflict in the latter, we speak consistently of taxonomic or phylogenomic *concepts*, emphasizing that these concepts are the immediate products of human empirical inference making [21].

Therefore, in aligning the neoavian explosion use case, we need not speak of the “same taxa” or “same clades” at all. Similarly, we need not judge whether one reconstruction or the other more closely aligns with deep-branching avian taxa, i.e., which is (more) ‘right’? Instead, our use case can fully develop within the context of modeling congruence and conflict across two sets of taxonomic *concept* hierarchies. We label these concepts with the “sec.” convention and maintain a one-to-one modeling relationship between concept labels and concepts. Accordingly, there is also no need to say that, in recognizing a concept with the taxonomic name Neornithes, the two teams are authoring “the same concept”. Instead, we model the two labels 2015.Neornithes and 2014. Neornithes, each of which symbolizes an individually generated phylogenomic theory region (concept). As an outcome of our alignment, we may say that these two concepts are *congruent*, or not, in a sense that reflects their relative alignment of the referential intension (to be specified below) of two phylogenomic theories. But by virtue of their differential source (author provenance), the two concepts 2015.Neornithes and 2014.Neornithes are never “the same”. In other words, “sameness” is rather trivially constrained in our representations to concepts whose labels contain an identical taxonomic name *and* which originate from a single phylogenomic hierarchy and source. That is, 2015.Neornithes and 2015.Neornithes are (labels for) the same concept.

### Knowledge representation and reasoning

The methods used herein are largely consistent with [14, 26, 32]. We refer to these publications for background and detail. Our logic representation utilizes three core conventions: (1) taxonomic concept labels to identify concepts; (2) *is_a* relationships to assemble single-source hierarchies via parent/child relationships; and (3) RCC–5 articulations to express the relative congruence of concept regions across multi-source hierarchies. The RCC–5 articulation vocabulary entails (with corresponding symbol): congruence (==), proper inclusion (>), inverse proper inclusion (<), overlap (><), and exclusion (!). Disjunctions of these articulations are a means to express uncertainty; as in: 2015.Neornithes {== *or* > *or* <} 2014.Neornithes. All possible disjunctions generate a lattice of 32 relationships (R_32_) that are expressible with RCC–5, where individual members of the “base five” are the most logically constraining subset [33].

The alignments are generated with the open source Euler/X software toolkit [28]. The toolkit can ingest multiple trees (T_1_, T_2_, etc.) and articulation sets (A), converting them into a set of logic constraints. Together with other default or facultative constraints (C) for modeling taxonomic hierarchies (see details below), these constraints are then submitted to a (suite of) logic reasoner(s) that achieve two main service tasks. First, the reasoner infers whether all input constraints are jointly logically consistent, i.e., whether they permit at least one “possible world”. Second, if consistency is attained, the reasoner infers the set of *Maximally Informative Relations* (MIR). The MIR constitute that unique set of RCC–5 articulations for every possible concept pair between the input sources from which the truth or falseness of any relationship in the R32 lattice can be deduced [14, 26, 33]. Many toolkit options and functions are designed to encode variable alignment input and output conditions, and to interactively obtain adequately constrained alignments. The toolkit also features a stylesheet-driven alignment input/output visualization service that utilizes directed acyclical graphs [28]. A step-wise account of the user/toolkit workflow interaction is provided in [26].

### Special challenges for multi-phylogeny alignments

Aligning phylogenomic trees entails several special representation and reasoning challenges. We address three aspects here that have not been dealt with extensively in previous publications.

#### 1. Representing intensional parent concept congruence via locally relaxed coverage

The first challenge relates directly to the notion of parent node identity in light of incongruently sampled child nodes. Unlike comprehensive classifications or revisions [14, 26, 34], phylogenomic reconstructions typically do not aspire to sample low-level entities exhaustively. Instead, select exemplars are sampled among all possible low-level entities, with the aim to represent phylogenomic diversity sufficiently well to infer reliable higher-level relationships. Often in practice, terminal sampling is not only incomplete for any single reconstruction, but purposefully complementary to that of other analyses. Generating informative genome-level data is resource-intensive [10]. This makes it prudent to coordinate terminal sampling globally, by prioritizing the reduction of gaps over redundant terminal sampling. In the case of 2015.PEA (198 terminals) versus 2014.JEA (48 terminals), only 12 species-level concept pairs have labels with identical taxonomic names.

By default, the Euler/X reasoning toolkit applies a *coverage constraint* to every input concept region. Coverage means that the region of a parent is strictly circumscribed by the union of its children [35]. However, this constraint is relaxable, either globally for all concepts, or locally for select concepts. The prefix “nc_” (no coverage), as in 2014.nc_Psittacidae, means: *either* a parent concept’s referential extension is circumscribed by the union of its explicitly included children, *or* there is a possibility of additional children being subsumed under the parent *but not mentioned* in the source phylogeny. Either scenario can yield consistent alignments. In other words, if a parent concept has relaxed coverage, it can potentially attain congruence with another parent concept in spite of each parent having a cumulatively incongruent set of child concepts. Trained judgment, reflected in the expert-asserted input articulations, can bring such instances of congruence to bear.

The desirable effect of locally relaxed coverage on aligning differentially sampled phylogenomic trees is illustrated in Figs. 1–4, using the example of parrots – 2015./2014.Psittaciformes. For this particular region, the author teams sampled wholly exclusively sets of concepts at the species level (Figs. 1 and 3). Even at the genus level, only 2015./2014.Nestor is redundantly sampled, yet with the articulation: 2015.Nestor_meridionalis ! 2014.Nestor_notabilis. Therefore, if no species-level concept sec. 2015.PEA has an explicitly sampled and congruent region in 2014.Psittaciformes, and vice-versa, if no species-level concept sec. 2014.JEA has such a region in 2015.Psittaciformes, then under global application of the coverage constraint we obtain the alignment: 2015.Psittaciformes ! 2014.Psittaciformes (Fig. 2). The absence of even partial concept region overlap at the terminal level ‘propagates up’ to the highest-level parent concepts, which are thereby also exclusive of each other.

**Fig 1.**
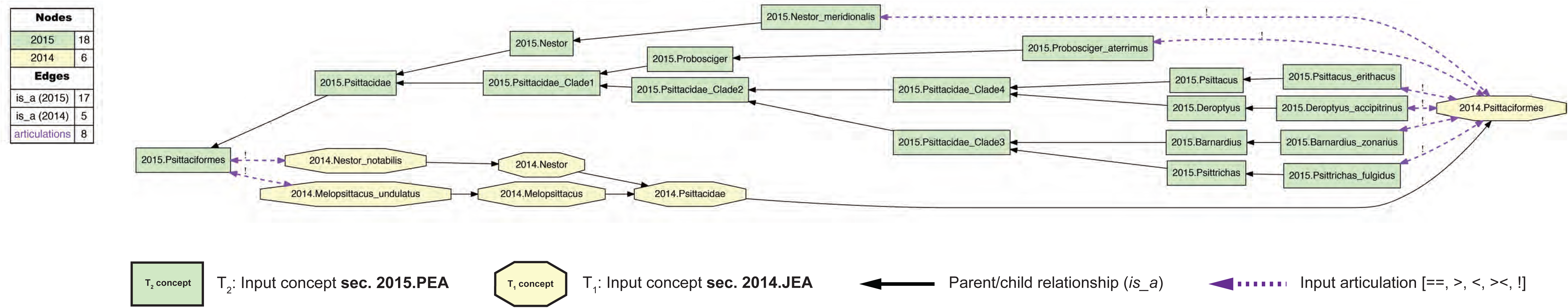
Input visualization for the 2015./2014.Psittaciformes alignment, with coverage globally applied. In all toolkit visualizations, the input and aligned, non-congruent concepts sec. 2015.PEA are shown as green rectangles (T_2_ – 18 concepts). Input and aligned, non-congruent concepts sec. 2014. JEA are shown as yellow octagons (T_1_ – 6 concepts). Congruent sets of aligned, multi-sourced concepts (first shown in Fig. 4) are rendered in gray rectangles with rounded corners. In this input visualization, each phylogenomic tree is separately assembled via parent/child (*is_a*) relationships (solid black arrows). All species-level concepts sec. 2015.PEA and 2014.JEA are exclusive of each other. Under strict application of the coverage constraint, this represented here by asserting eight articulations (dashed magenta arrows) of disjointness (!) of each species-level concept from the other-sourced order-level concept. The legend indicates the number of nodes and edges for each input tree, and the number of parent/child relationships and expert-asserted input articulations. See also S1A and S1B Files.

**Fig 2.**
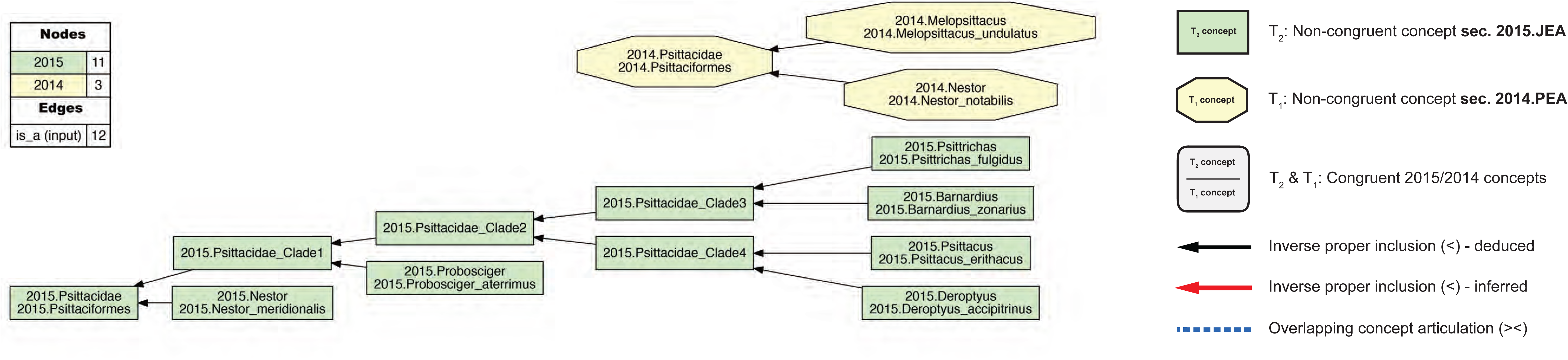
Alignment visualization for the 2015./2014.Psittaciformes alignment, with coverage globally applied. This alignment corresponds to the Fig. 1 input, and shows reasoner-inferred non-/congruent concepts and articulations (see legend) – i.e., none in this particular case. The reasoner infers 108 logically implied articulations that constitute the set of MIR. See also S2A and S2B Files.

**Fig 3.**
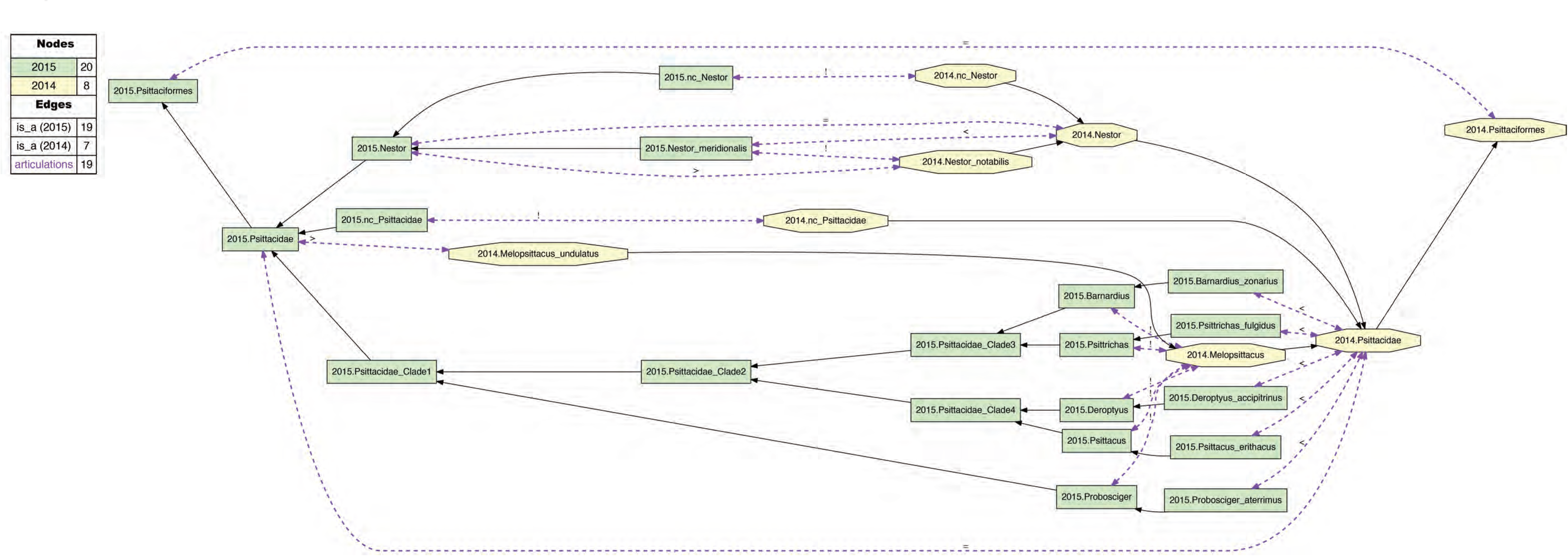
Input visualization for the 2015./2014.Psittaciformes alignment, with coverage locally relaxed. Compare with Fig. 1. Here, coverage is relaxed for two family-level concepts (2015./2014.nc_Pittacidae) and two genus-level concepts (2015./2014.nc_Nestor). The eight species-level concepts of the alignment are correspondingly included as members of these higher-level concepts. In addition, three instances of congruence are asserted for 2015. /2014.{Psittaciformes, Psittacidae, Nestor}. See also S3A and S3B Files.

**Fig 4.**
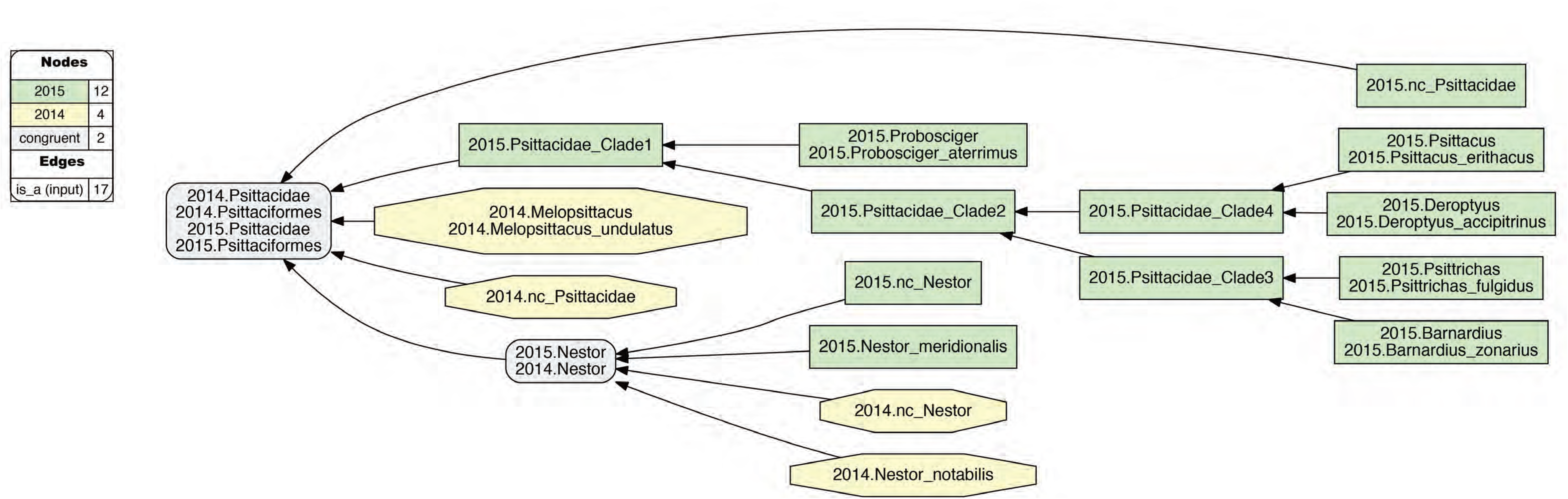
Alignment visualization for the 2015./2014.Psittaciformes alignment, with coverage locally relaxed. Compare with Fig. 2. Local relaxing of coverage and assertions of congruence of paired higher-level concepts (Fig. 3) yields the intuitive alignment of 2015.Psittaciformes == 2014. Psittaciformes, 2015.Psittacidae == 2014.Psittacidae, and 2015.Nestor = 2014.Nestor; in spite of wholly incongruent sampling of species-level concepts. The reasoner infers 160 logically implied articulations that constitute the set of MIR. See also S4A and S4B Files.

Although the Figs. 1 and 2 input and alignment are empirically defensible and logically consistent, they fail to capture certain intuitions we have regarding the higher-level 2015./2014.Psittaciformes relationship. For instance, we may wish to say: “Sure, the author teams sampled complementary species-level concepts. Yet these trees are not actually in conflict. At higher levels, there likely is agreement that parrots are parrots, and non-parrots are non-parrots”. That is: 2015.Psittaciformes == 2014.Psittaciformes. To obtain this intuitive alignment, we have to *locally* relax coverage at select lower levels (Fig. 3). In particular, 2015.PEA include five genus- and species-level concepts under 2015.Psittacidae that have no corresponding region under Psittacidae. However, if we relax coverage for 2014.Psittacidae – i.e., we assert 2014. nc_Psittacidae as an input concept or constraint – then we can include each of these; for instance: 2015.Probosciger_aterrimus < 2014.Psittacidae, 2015.Psittacus_erithacus < 2014. Psittacidae, etc. Conversely, if we locally relax coverage for 2015.Psittacidae (2015.nc_Psittacidae), we can specify 2014.Melopsittacus_undulatus < 2015.Psittacidae. At the genus level, we can align 2015.Nestor == 2014.Nestor if we relax coverage for each (2015.nc_Nestor, 2014.nc_Nestor), in spite of the mutually exclusive species-level concepts sampled (see above). Jointly, these four instances of relaxing coverage render the articulation 2015. Psittacidae == 2014.Psittacidae consistent, and hence also 2015.Psittaciformes == 2014. Psittaciformes (Fig. 4).

Expressing higher-level node congruence in light of lower-level node incongruence requires a conception of node identity that affirms counter-factual statements of the following type: if 2014. JEA had sampled 2014.Psittacus_erithacus, then the authors would have included this species-level concept as a child of 2014.Psittacidae. This is to say that 2015./2014.Psittacidae, and hence their respective parents, are *intensionally* defined [25, 36, 37]. Using trained judgment [30], we align these concept regions as if there are congruent *property* criteria that each region entails, i. e., something akin to an implicit set of synapomorphies or uniquely diagnostic features. Of course, the phylogenomic data provided by 2015.PEA and 2014.JEA do not signal intensional definitions directly. But neither do their genome-based topologies for parrots provide evidence to challenge the status of such definitions as previously proposed [38]. Unlike ostensive parent node definitions, which would point at one or more exemplary children, or extensional definitions, which would specify all children exhaustively, intensional definitions have predictive powers regarding the inclusion of children under a parent concept which does not explicitly list them (all). Therefore, given the incongruent low-level node sampling, asserting reciprocally congruent intensional parent node definitions is the only way to obtain 2015.Psittacidae == 2014.Psittacidae, or 2015.Psittaciformes == 2014.Psittaciformes.

A clarification is in place. Region Connection Calculus is at best a means of *translating* the signal of an intensional definition. The congruent (==) symbol means, only: two regions are congruent in their extension. The RCC–5 vocabulary is obviously not appropriate for reasoning directly over genomic or phenomic property statements. The reasoner does not assess whether 2015.Psittacidae, or any included child or aligned concept, has ‘the relevant synapomorphies’. Doing so would not be trivial even if property-based definitions were provided, because we would still have to make theory-laden assumptions about their congruent phylogenomic scopes [26, 39, 40].

In summary, locally relaxing coverage is the only means under the RCC–5 reasoning approach to obtain intuitive, logically consistent and congruent higher-level concept alignments when lower-level sampling is incongruent. We *interpret* this practice as analogous to defining parent concept regions intensionally, even though the source publications do not provide explicit intensional node definitions, and in any case, the language of RCC–5 can at most provide a signal translation of a property-based definition. Nevertheless, we will see below that locally relaxing coverage is an essential tool for exposing meaningful higher-level node identities and relationships in the neoavian explosion use case.

#### 2. Representing clade concept labels

Our modeling approach requires that every region in each source tree receives a taxonomic or clade concept label. However, the source publications only provide such labels for a subset of the inferred nodes. In particular, 2015.PEA (p. 570: figure 1) obtained 41 nodes above the ordinal level. Of these, 17 nodes (41.5%) were explicitly labeled in either the published figure or supplement (pp. 9-12). The authors also cite [20] as the primary source for valid name usages, yet this list is not concerned with supra-ordinal names. Similarly, 2014. JEA (p. 1322: figure 1) inferred 37 nodes above the ordinal level, of which 23 nodes (62.2%) were given an explicit label. They provide an account (cf. supplementary materials SM6: 22-24) of their preferred name usages, sourced mainly to [20] and [41].

In assigning clade concept labels at the supra-ordinal level when the authors may have failed to do so (consistently), we nevertheless made a good faith effort – through examination of the supplementary information and additional sources [1, 3, 42, 43, 44, 45, 46, 47] – to represent the authors’ preferred name usages. Where usages were not explicit, we selected the only or most commonly applied clade concept name at the time of publication of the phylogenomic reconstructions. This effort yielded 13 additional labels for 2015.PEA (Table 1), and 7 such labels for 2014.JEA (Table 2).

**Table 1.** Supra-ordinal clade concept labels used for the phylogenomic tree of 2015.PEA, with sources from which the names were obtained. “Franz et al. 2017” means: the label was assigned pragmatically in this study. See main text for further detail.

| # | Clade concept label | Utilized name source | Immediate child concepts |
| --- | --- | --- | --- |
| P01 | 2015.Neornithes | Livezey & Zusi 2007 [43] | 2015.Palaeognathae, 2015.Neognathae |
| P02 | 2015.Palaeognathae | Prum et al. 2015 [2] | 2015.Notopalaeognathae, 2015.Struthioniformes |
| P03 | 2015.Notopalaeognathae | Yuri et al. 2013 [46] | 2015.Novaeratitae, 2015.Rheiformes |
| P04 | 2015.Novaeratitae | Yuri et al. 2013 [46] | 2015.Apterygiformes, 2015.Novaeratitae_Clade1 |
| P05 | 2015.Novaeratitae_Clade1 | Franz et al. 2017 | 2015.Casuariiiformes, 2015.Tinamiformes |
| P06 | 2015.Neognathae | Jarvis et al. 2014 [1] | 2015.Galloanserae, 2015.Neoaves |
| P07 | 2015.Galloanserae | Prum et al. 2015 [2] | 2015.Anseriiformes, 2015.Galliiformes |
| P08 | 2015.Neoaves | Prum et al. 2015 [2] | 2015.Strisores, 2015.Neoaves_Clade1 |
| P09 | 2015.Strisores | Prum et al. 2015 [2] | 2015.Caprimulgidae, 2015.Strisores_Clade1 |
| P10 | 2015.Strisores_Clade1 | Franz et al. 2017 | 2015.Nyctibiidae, 2015.Steatornithidae, 2015.Strisores_Clade2 |
| P11 | 2015.Strisores_Clade2 | Franz et al. 2017 | 2015.Apodiformes, 2015.Aegothelidae, 2015.Podargidae |
| P12 | 2015.Neoaves_Clade1 | Franz et al. 2017 | 2015.Columbaves, 2015.Neoaves_Clade2 |
| P13 | 2015.Columbaves | Prum et al. 2015 [2] | 2015.Columbimorphae, 2015.Otidimorphae |
| P14 | 2015.Columbimorphae | Prum et al. 2015 [2] | 2015.Columbiformes, 2015.Columbimorphae_Clade1 |
| P15 | 2015.Columbimorphae_Clade1 | Franz et al. 2017 | 2015.Mesitornithiformes, 2015.Pterocliiformes |
| P16 | 2015.Otidimorphae | Prum et al. 2015 [2] | 2015.Musophagiformes, 2015.Otidimorphae_Clade1 |
| P17 | 2015.Otidimorphae_Clade1 | Franz et al. 2017 | 2015.Mesitornithiformes, 2015.Pteroclidiformes |
| P18 | 2015.Neoaves_Clade2 | Franz et al. 2017 | 2015.Gruiformes, 2015.Neoaves_Clade3 |
| P19 | 2015.Neoaves_Clade3 | Franz et al. 2017 | 2015.Aequorlornithes, 2015.Inopinaves |
| P20 | 2015.Aequorlornithes | Prum et al. 2015 [2] | 2015.Aequorlornithes_Clade1, 2015.Ardeae |
| P21 | 2015.Aequorlornithes_Clade1 | Franz et al. 2017 | 2015.Charadriiformes, 2015.2015.Phoenicopterimorphae |
| P22 | 2015.Phoenicopterimorphae | Jarvis et al. 2014 [1] | 2015.Phoenicopteriformes, 2015.Podicipediformes |
| P23 | 2015.Ardeae | Brodkorb 1963 [42] | 2015.Aequornithia, 2015.Phaethontimorphae |
| P24 | 2015.Aequornithia | Prum et al. 2015 [2] | 2015.Aequornithia_Clade1, 2015.Gaviiformes |
| P25 | 2015.Aequornithia_Clade1 | Franz et al. 2017 | 2015.Pelecanimorphae, 2015.Procellariimorphae |
| P26 | 2015.Pelecanimorphae | Prum et al. 2015 [2] | 2015.Ciconiiformes, 2015.Pelecanimorphae_Clade1 |
| P27 | 2015.Pelecanimorphae_Clade1 | Franz et al. 2017 | 2015.Pelecaniformes, 2015.Suliformes |
| P28 | 2015.Procellariimorphae | Prum et al. 2015 [2] | 2015.Procellariiformes, 2015.Sphenisciformes |
| P29 | 2015.Phaethontimorphae | Prum et al. 2015 [2] | 2015.Eurypygiformes, 2015.Phaethontiformes |
| P30 | 2015.Inopinaves | Prum et al. 2015 [2] | 2015.Opisthocomiformes, 2015.Telluraves |
| P31 | 2015.Telluraves | Prum et al. 2015 [2] | 2015.Accipitriformes, 2015.Eutelluraves |
| P32 | 2015.Eutelluraves | Prum et al. 2015 [2] | 2015.Australaves, 2015.Coracornithia |
| P33 | 2015.Australaves | Prum et al. 2015 [2] | 2015.Cariamiformes, 2015.Eufalconimorphae |
| P34 | 2015.Eufalconimorphae | Suh et al. 2011 [45] | 2015.Falconiformes, 2015.Passerimorphae |
| P35 | 2015.Passerimorphae | Sibley et al. 1988 [3] | 2015.Passeriformes, 2015.Psittaciformes |
| P36 | 2015.Coracornithia | Claramunt & Cracraft 2015 [47] | 2015.Coraciimorphae, 2015.Strigiformes |
| P37 | 2015.Coraciimorphae | Prum et al. 2015 [2] | 2015.Coliiformes, 2015.Eucavitaves |
| P38 | 2015.Eucavitaves | Yuri et al. 2013 [46] | 2015.Cavitaves, 2015.Leptosomiformes |
| P39 | 2015.Cavitaves | Yuri et al. 2013 [46] | 2015.Picocoraciae, 2015.Trogoniformes |
| P40 | 2015.Picocoraciae | Mayr 2010 [44] | 2015.Bucerotiformes, 2015.Picodynastornithes |
| P41 | 2015.Picodynastornithes | Yuri et al. 2013 [46] | 2015.Coraciiformes, 2015.Piciformes |

**Table 2.** Supra-ordinal clade concept labels used for the phylogenomic tree of 2014.JEA, with sources from which the names were obtained. “Franz et al. 2017” means: the label was assigned pragmatically in this study. See main text for further detail.

| # | 2014.JEA clade concept label | Utilized name source | Immediate child concepts |
| --- | --- | --- | --- |
| J01 | 2014.Neornithes | Jarvis et al. 2014 [1] | 2014.Palaeognathae, 2014.Neognathae |
| J02 | 2014.Palaeognathae | Jarvis et al. 2014 [1] | 2014.Struthioniformes, 2014.Tinamiformes |
| J03 | 2014.Neognathae | Jarvis et al. 2014 [1] | 2014.Galloanseres, 2014.Neoaves |
| J04 | 2014.Galloanseres | Jarvis et al. 2014 [1] | 2014.Anseriiformes, 2014.Galliiformes |
| J05 | 2014.Neoaves | Jarvis et al. 2014 [1] | 2014.Columbea, 2014.Passerea |
| J06 | 2014.Columbea | Jarvis et al. 2014 [1] | 2014.Columbimorphae, 2014.Phoenicopterimorphae |
| J07 | 2014.Columbimorphae | Jarvis et al. 2014 [1] | 2014.Columbiformes, 2014.Columbimorphae_Clade1 |
| **J08** | 2014.Columbimorphae_Clade1 | Franz et al. 2017 | 2014.Mesitornithiformes, 2014.Pterocliiformes |
| **J09** | 2014.Phoenicopterimorphae | Jarvis et al. 2014 [1] | 2014.Phoenicopteriformes, 2014.Podicipediformes |
| **J10** | 2014.Passerea | Jarvis et al. 2014 [1] | 2014.Passerea_Clade1, 2014.Passerea_Clade4 |
| **J11** | 2014.Passerea_Clade1 | Franz et al. 2017 | 2014.Passerea_Clade2, 2014.Passerea_Clade3 |
| **J12** | 2014.Passerea_Clade2 | Franz et al. 2017 | 2014.Ardeae, 2014.Telluraves |
| **J13** | 2014.Ardeae | Brodkorb 1963 [42] | 2014.Aequornithia, 2014.Phaethontimorphae |
| **J14** | 2014.Aequornithia | Jarvis et al. 2014 [1] | 2014.Aequornithia_Clade1, 2014.Gaviimorphae |
| **J15** | 2014.Aequornithia_Clade1 | Franz et al. 2017 | 2014.Pelecanimorphae, 2014.Procellariimorphae |
| **J16** | 2014.Pelecanimorphae | Jarvis et al. 2014 [1] | 2014.Pelecaniformes |
| **J17** | 2014.Procellariimorphae | Jarvis et al. 2014 [1] | 2014.Procellariiformes, 2014.Sphenisciformes |
| **J18** | 2014.Gaviimorphae | Jarvis et al. 2014 [1] | 2014.Gaviiformes |
| **J19** | 2014.Phaethontimorphae | Jarvis et al. 2014 [1] | 2014.Eurypygiiformes, 2014.Phaethontiformes |
| **J20** | 2014.Telluraves | Jarvis et al. 2014 [1] | 2014.Afroaves, 2014.Australaves |
| **J21** | 2014.Afroaves | Jarvis et al. 2014 [1] | 2014.Accipitrimorphae, 2014.Coracornithia |
| **J22** | 2014.Accipitrimorphae | Jarvis et al. 2014 [1] | 2014.Accipitriiformes |
| **J23** | 2014.Coracornithia | Claramunt & Cracraft 2015 [47] | 2014.Coraciimorphae, 2014.Strigiformes |
| **J24** | 2014.Coraciimorphae | Jarvis et al. 2014 [1] | 2014.Coliiformes, 2014.Eucavitaves |
| **J25** | 2014.Eucavitaves | Yuri et al. 2013 [46] | 2014.Cavitates, 2014.Leptosomiformes |
| **J26** | 2014.Cavitates | Yuri et al. 2013 [46] | 2014.Picocoraciae, 2014.Trogoniformes |
| **J27** | 2014.Picocoraciae | Mayr 2010 [44] | 2014.Bucerotiformes, 2014.Picodynastornithes |
| **J28** | 2014.Picodynastornithes | Yuri et al. 2013 [46] | 2014.Coraciiformes, 2014.Piciformes |
| **J29** | 2014.Australaves | Jarvis et al. 2014 [1] | 2014.Cariamiformes, 2014.Eufalconimorphae |
| **J30** | 2014.Eufalconimorphae | Suh et al. 2011 [45] | 2014.Falconiformes, 2014.Passerimorphae |
| **J31** | 2014.Passerimorphae | Jarvis et al. 2014 [1] | 2014.Passeriformes, 2014.Psittaciformes |
| **J32** | 2014.Passerea_Clade3 | Franz et al. 2017 | 2014.Cursorimorphae, 2014.Opisthocomiformes |
| **J33** | 2014.Cursorimorphae | Jarvis et al. 2014 [1] | 2014.Charadriiformes, 2014.Gruiformes |
| **J34** | 2014.Passerea_Clade4 | Franz et al. 2017 | 2014.Caprimulgimorphae, 2014.Otidimorphae |
| **J35** | 2014.Caprimulgimorphae | Jarvis et al. 2014 [1] | 2014.Caprimulgiformes |
| **J36** | 2014.Otidimorphae | Jarvis et al. 2014 [1] | 2014.Cuculiformes, 2014.Otidimorphae_Clade1 |
| **J37** | 2014.Otidimorphae_Clade1 | Franz et al. 2017 | 2014.Musophagiformes, 2014.Otidiformes |

If no suitable label was available, we chose a simple naming convention of adding “_Clade1”, “_Clade2”, etc., to the available and immediately higher-level node label, e.g. 2014. Passerea_Clade1. The numbering of such labels along the tree topology starts with the most immediate child of a properly named parent, and typically follows down one section of the source tree entirely before continuing with the higher-level sister section. Using this approach, we added 11 labels for 2015.PEA (Table 1) and 7 labels for JEA.2014 (Table 2).

The clade concept labeling convention was not applied below the family level, where instead we opted to collapse phylogenomic resolution such that all genus-level concepts are the immediate (polytomous) children of the parent (exception: Figs. 1–4). This was done because in the case of 2014. JEA, only four family-level concepts include two children, whereas the remainder have a single child sampled. Clearly, resolving the monophyly of subfamilial clade concepts was not the primary aim of 2014.JEA. The same applies to 2015.PEA, who sampled 104/125 family-level concepts with only 1-2 children. Thus, a pragmatic choice was made to simplify the alignments, by not representing subfamilial clade concepts in the 21 instances provided by 2015.PEA.

#### 3. Representing phylogeny/classification paraphyly

A third, relatively minor challenge is the occurrence of clade concepts in 2015.PEA’s phylogenomic tree that are not congruently aligned with higher-level concepts of [20]. We highlight these instances here because they represent a widespread phenomenon in phylogenomics. It is useful to understand how such discrepancies can be modeled with RCC–5 alignments (Figs. 5 and 6).

**Fig 5.**
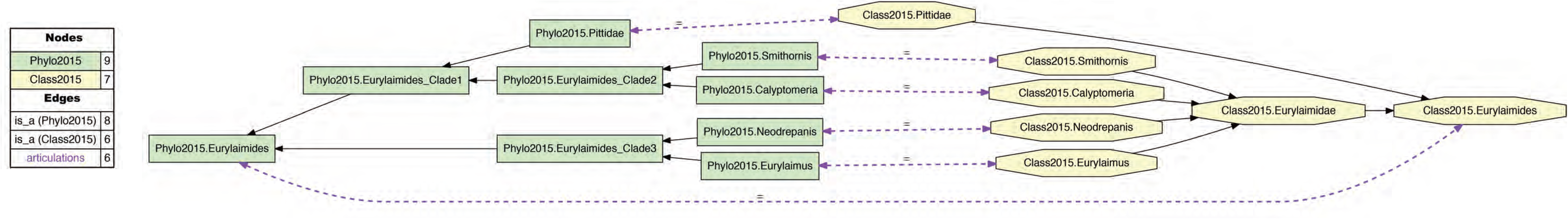
Input visualization of the alignment of the phylogenomic reconstruction of passeriform *clade* concepts sec. 2015.PEA – prefixed with “Phylo2015” – with the corresponding *classification* concepts sec. Gill & Donsker (2015) [20] – prefixed with “Class2015”. The phylogenomic topology renders that of Class2015.Eurylaimidae paraphyletic, and hence the name “Eurylaimidae” is not represented in any clade concept label sec. 2015.PEA. See also Prum et al. (2015). See also S5A and S5B Files.

**Fig 6.**
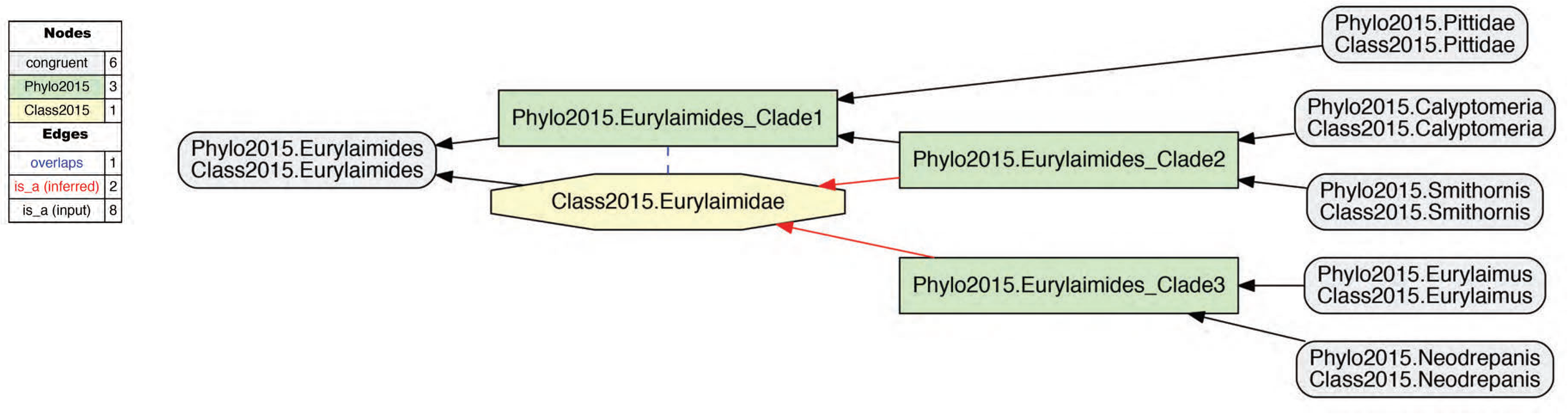
Alignment visualization corresponding to Fig. 5. The alignment shows an overlapping articulation (dashed blue line) between the phylogenomic clade concept sec. 2015.PEA (Phylo2015.Passeriformes_Clade1) and the Eurylaimidae sec. Gill & Donsker (2015) [20] (Class2015.Eurylaimidae). The two dashed red arrows symbolized reasoner-inferred relationships not explicit in the input constraints. See also S6A and S6B Files.

Figure 5 exemplifies the phylogenetic tree/classification incongruence observed in 2015.PEA. The authors state (supplementary table 1, p. 1): “Taxonomy follows Gill and Donsker (2015; fifth ed)”. As shown in Fig. 5, their phylogeny accommodates four sampled genus-level concepts that would correspond to children of the family-level concept Eurylaimidae sec. Gill & Donsker (2015) [20]. However, these concepts are arranged paraphyletically in relation to the reference classification. This mean that there is no parent concept in 2015.PEA’s reconstruction that can be labeled 2015.Eurylaimidae that would not also (1) include 2015.Pittidae, i.e., 2015.Passeriformes_Clade1 in Figure 6, or (2) just represent aligned subset of the Eurylaimidae sec. Gill and Donsker (2015) [20], i.e., 2015.Passeriformes_Clade2 or 2015.Passeriformes_Clade3 in Figure 6. The concept Eurylaimidae sec. Gill and Donsker (2015) [20] has an overlapping (><) articulation with 2015.Passeriformes_Clade1.

In summary, our approach represents paraphyly (or any kind of non-monophyly) as an incongruent alignment of the phylogenomic tree and the source classification used – though not logically suited – to provide labels for that tree’s monophyletic clade concepts. There are four distinct regions in the phylogeny of 2015.PEA where such alignments are needed to represent incongruence with these taxonomic concept labels: {Caprimulgiformes, Eurylaimidae, Hydrobatidae, Procellariidae, Tityridae} sec. Gill & Donsker (2015) [20]. Each of these is provided in the S7-S9 File sets, which provide an understanding of the phylogeny/classification incongruence internal to 2015.PEA’ labeling conventions. For the purpose of aligning the 2015./2014.Neornithes phylogenies, we adhere to the clade concept labels of Tables 1 and 2.

### Configuration of input constraints and alignment partitioning

The size and complexity of the use case present certain logic reasoning and visualization constraints. As published, the source phylogenies specify 703 and 216 clade or taxonomic concepts, respectively. The use case is therefore comparable in size to [14]. However, the frequent instances of locally relaxed coverage increase the reasoning complexity, to the point where custom RCC–5 reasoning is needed [48]. The reasoning challenges, together with the difficulty of visualizing nearly 920 concept labels legibly in publication format, commend a partitioned alignment approach. To keep the Results concise, we show visualizations of the larger input and alignment partitions only in the Supporting Information. A detailed account of the input configuration and partitioning workflow is given below.

For complex alignment challenges, the toolkit workflow favors a partitioned, bottom-up approach [29]. The large problem of aligning all concepts is broken down into multiple smaller alignment problems, e.g. 2015./2014.Psittaciformes (Figs. 1–4). In each case, RCC–5 articulations for low-level concept pairs are provided incrementally, e.g., in sets of 1-5 articulations at a time. Following such an increment, the toolkit reasoning process is re-/deployed to validate input consistency and infer the number of possible worlds. This stepwise approach leads to increasingly constrained alignments, along with user-manageable products [26]. Once a set of small, topographically adjacent alignment partitions is well specified, these can serve as building blocks for the next, larger partition. An example of the latter is the 2015./2014.Passerimorphae alignment, which includes two order-level concepts and their children in each source phylogeny. Such midlevel partitions eventually form the basis for the largest alignment partitions, e.g. 2015./2014.Telluraves.

Underlying all alignments is the presumption that at the terminal (species) level, the taxonomic concept labels of 2015.PEA and 2014.JEA are reliable indicators of either pairwise congruence or exclusion [14, 26, 32]. That is, e.g., 2015.Cariama_cristata == 2014.Cariama_cristata, or 2015.Charadrius_hiaticula ! 2014.Charadrius_vociferus. Because the time interval separating the two publications is short in comparison to the time needed for taxonomic revisions to effect changes in classificatory practice, the genus- or species-level taxonomic concepts are unlikely to show much incongruence; though see [49] or [50]. We note that 2015.PEA (p. 571) use the label 2015.Urocolius(_indicus) in their phylogenomic tree, which also corresponds to the genus-level name endorsed in [20] Gill & Donsker (2015). However, in their Supplementary Table 1 the authors use 2015.Colius_indicus. We chose 2015.Urocolius and 2015.Urocolius_indicus as the labels to apply in the alignments.

In all, 2014.JEA sample children of 34 order-level concepts in their phylogeny, whereas 2015.PEA recognize 40 order-level concepts. The latter authors represent four order-level concepts for which no analogous children are included in 2014.JEA, i.e.: 2015.{Apterygiformes, Casuariiformes, Ciconiiformes, Rheiformes}. Three of these are assigned to 2015.Palaeognathae, whereas 2015.Ciconiiformes are subsumed under 2015.Pelecanimorphae. The remaining 36 order-level concepts sec. 2015.PEA show some child-level overlap with those of 2014.JEA. Thus, a suitable partitioning approach starts with specifying the input constraints for nearly 35 paired order-level concepts and their respective children, as demonstrated in Figs. 3 and 4. The largest order-level partition is 2015./2014.Passeriformes, with 148 x 22 input concepts, seven instances of relaxed parent coverage, and 101 input articulations. This alignment completes in less than 15 seconds on an individual 2.0 GHz processor, yielding 3,256 MIR.

Once an order-level partition is completed, the reasoner-inferred higher-level concept relationships can be added as articulations to the set of input constraints for that alignment. This is a means of further constraining how the alignment will 'behave' in an expanded context, i.e., how many possible worlds the combination of multiple sub-partitions into one larger partition may permit. In most cases, given the use of the “no coverage” approach illustrated above, congruence is attained at the ordinal concept level. These completed partitions can therefore act as congruent 'terminals' for consistent, supra-ordinal alignments.

We configured six larger, non-overlapping partitions as building blocks for the global alignment: 2015./2014.Palaeognathae (34 x 12 input concepts, four instances of relaxed coverage, and 25 articulations; same data sequence used for following alignments), 2015./2014.Galloanserae (49, 16, 7, 46), 2015.Columbaves/2014.Columbimorphae + 2014.ütidimorphae (53, 37, 13, 37), 2015.Strisores/2014.Caprimulgimorphae (44, 17, 8, 32), 2015./2014.Ardeae (100, 55, 19, 75), and the largest partition of 2015./2014.Telluraves (316, 104, 37, 241).

The inferred congruence of 2015.Telluraves == 2014.Telluraves presents an opportunity to partition the entire alignment into two similarly sized regions, where the complementary region includes all 2015./2014.Neornithes concepts (392, 174, 58, 259), *except* those subsumed under 2015./2014.Telluraves, which are therein only represented with two concepts labels and one congruent articulation. These two complements – i.e., 2015./2014.Neornithes (without) / 2015./2014.Telluraves – are the core partitions that inform our use case alignment, globally. The corresponding S10-S11 File sets include the input constraint (.txt) and visualization (.pdf) files, along with the alignment visualization (.pdf) and MIR (.csv).

The two large partitions yield unambiguous RCC–5 articulations from the species concept level to that of 2015./2014.Neornithes. They can be aggregated into a synthetic, root-to-order level alignment, where all subordinal concepts and articulations are secondarily pruned away. Such an alignment retains the logic signal derived from the bottom-up approach, but represents only congruent order-level concept labels as terminal regions, except in cases where there is incongruence (e.g., 2015.Pelecaniformes < 2014.Pelecaniformes; see Introduction). We present this alignment as an analogue to figure 1 in [4] (p. 515), and compare how each conveys information about congruent and conflicting clade concepts.

Lastly, we further reduce the root-to-order alignment to display only 5-6 clade concept levels below the congruent 2015./2014.Neoaves. This region of the alignment is the most conflicting; hence, modeling this conflict with adequate granularity forms the basis for our Discussion.

## Results

### Higher-level congruence

Our alignments show widespread higher-level congruence across the neoavian explosion use case; along with several minor regions of conflict and one strongly conflicting region – as expected – between concepts placed immediately below the 2015./2014.Neoaves.

We focus first on the two large complementary alignment partitions, i.e. 2015./2014.Neornithes (without) / 2015./2014.Telluraves (see S10 and S11 File sets). Jointly, they entail 707 concepts sec. 2015.PEA and 283 concepts sec. 2014.JEA. Among these, 34 “no coverage” regions were added to 2015.PEA’s phylogeny, whereas 61 instances of relaxing parent coverage were assigned to 2014.JEA’s phylogeny. The 2015./2014.Neornithes partition shows 305 aligned regions – 247 without the “no coverage” regions – of which 60 congruently carry at least one concept label from each source phylogeny. This alignment also shows eight congruent species-level concept regions; the latter would be the only instances of congruence if coverage were globally applied (Figs. 1 and 2). Therefore, relaxing the coverage constraint yields 52 *additional* instances of higher-level node congruence. Similarly, the 2015./2014.Telluraves partition has 231 aligned regions – 194 without the “no coverage” regions – of which 38 are congruent. This corresponds to an increase of 34 regions, compared to the four congruent species-level concept regions present under strict coverage. Correcting for the redundant 2015./2014.Telluraves region, we ‘gain’ 85 congruent parent node regions across the two phylogenies *if* node identity is encoded intensionally (Figs. 3 and 4). Indeed, this approach yields the logically consistent, intuitive articulation 2015.Neornithes == 2014.Neornithes at the highest level. We might say: “Regardless of sampling biases, the two author teams concur that modern birds are modern birds”.

### Two kinds of conflict: Differential granularity and overlap

We now focus on characterizing topological conflict between 2015.PEA and 2014.JEA. Phylogenomic incongruence can be divided into two general categories: (1) differential granularity or resolution of clade concepts (RCC–5 translation: < or >), and (2) overlapping clade concepts (RCC–5 translation: ><).

The first of these is less problematic from a standpoint of achieving data integration. Given a particular subregion of the alignment, the more densely sampled phylogeny will entail additional, more finely resolved clade concepts in comparison to its counterpart. Typically, this distinction belongs to the phylogeny of 2015.PEA, due to the 4:1 ratio of terminals sampled. Indeed, there are 83 above species-level clade concepts sec. 2015.PEA that can be interpreted as *congruent refinements* of the 2014.JEA topology (see S10 and S11 File sets). Conversely, only two such instances of added resolution are contributed by 2014.JEA: (1) 2014.Passeriformes_Clade3 which entails 2014.Passeridae and 2014.Thraupidae; and (2) 2014.Haliaeetus with two subsumed species-level concepts. Nevertheless, the joint 97 congruent node regions and 85 refining node regions cover a large section of the alignment where data integration is either reciprocally (==) or unilaterally (< or >) feasible.

### Focus on topological overlap

The remaining 38 instances of overlapping articulations between higher-level 2015/2014 concepts constitute the most profound cases of conflict. These instances are clustered in four distinct regions, i.e.: 2015./2014.Pelecanimorphae (8 overlaps; Fig. 7 and S12 File set); 2015.Passeri/2014.Passeriformes_Clade2 (3 overlaps; Fig. 8 and S13 File set); 2015.Eutelluraves/2014.Afroaves (1 overlap; Figs. 9 and 10, and S14 and S15 File sets); and finally, 2015./2014.Neoaves (26 overlaps; Figs. 11–13, and S16-S18 File sets). We will examine each of these in sequence.

**Fig 7.**
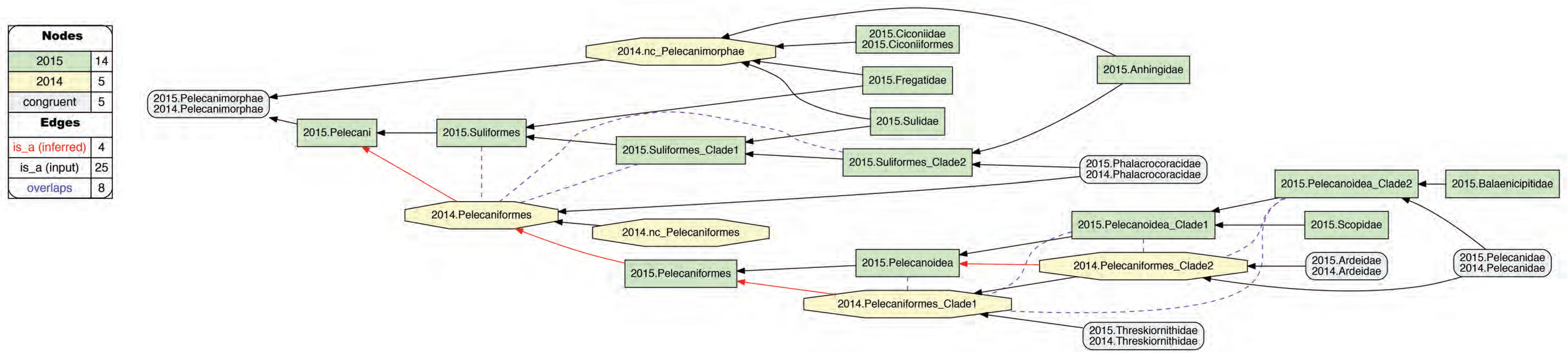
Alignment visualization for the 2015./2014.Pelecanimorphae alignment, with eight overlapping relationships. See text for further detail. The reasoner infers 200 logically implied articulations that constitute the set of MIR. See also S12 File set.

**Fig 8.**
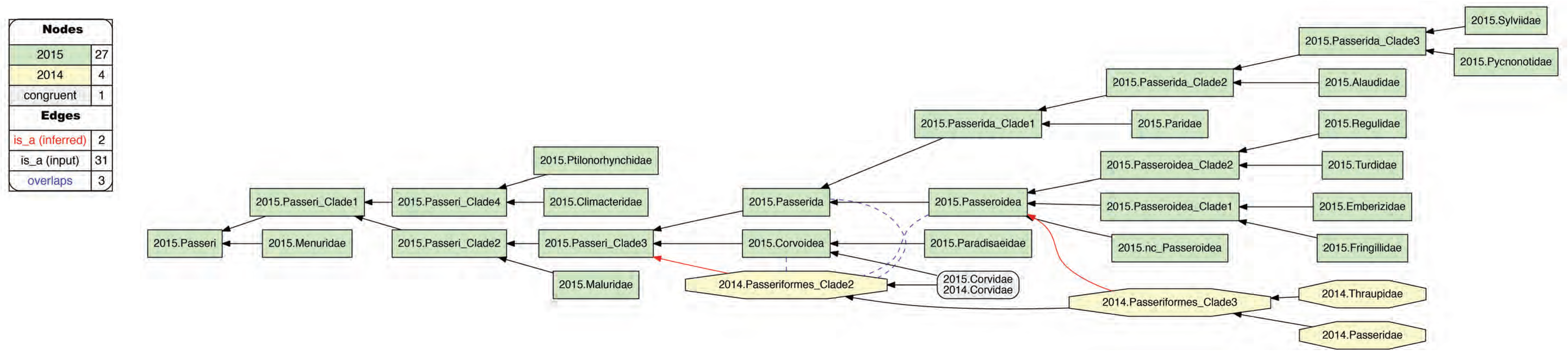
Alignment visualization for the 2015.Passeri/2014.Passeriformes_Clade2 alignment, with three overlapping relationships. See text for further detail. The reasoner infers 135 logically implied articulations that constitute the set of MIR. See also S13 File set.

**Fig 9.**
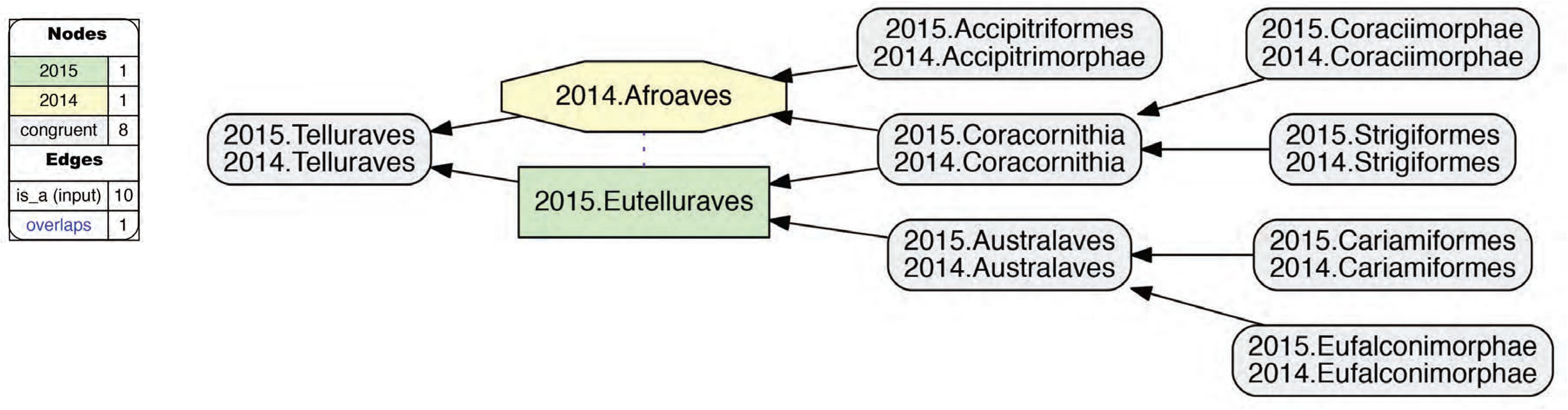
Alignment visualization for the 2015./2014.Telluraves alignment, under whole-concept resolution, with one overlapping relationship. Compare with Fig. 10; see text for further detail. The reasoner infers 81 logically implied articulations that constitute the set of MIR. See also S14 File set.

**Fig 10.**
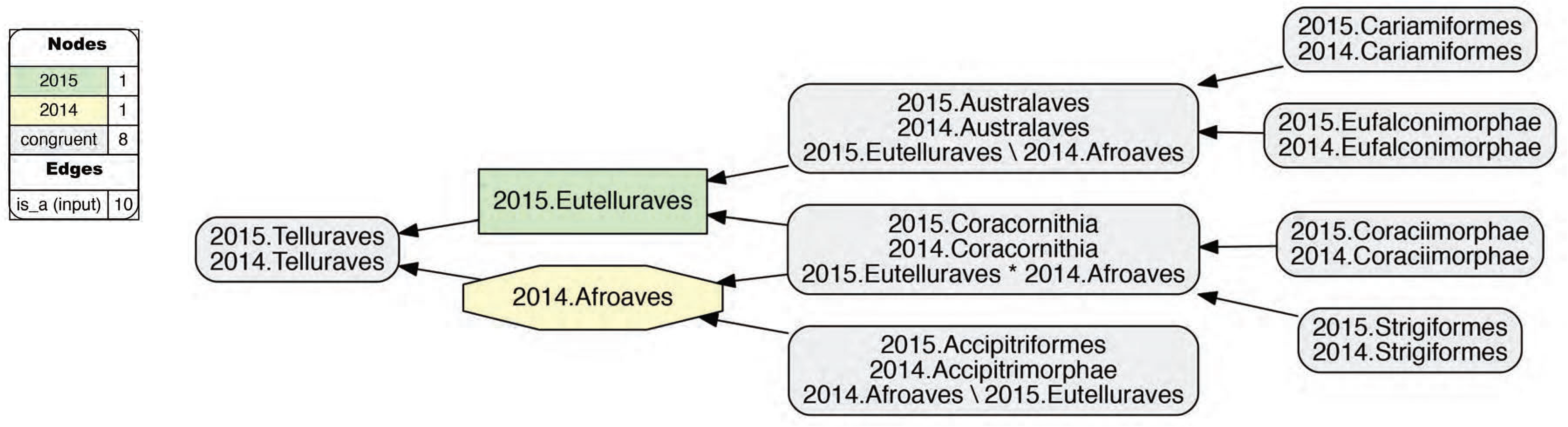
Alignment visualization for the 2015./2014.Telluraves alignment, under split-concept resolution, resolving the overlapping relationship. Compare with Fig. 9; see text for further detail. The reasoner infers 81 logically implied articulations that constitute the set of MIR. See also S15 File set.

**Fig 11.**
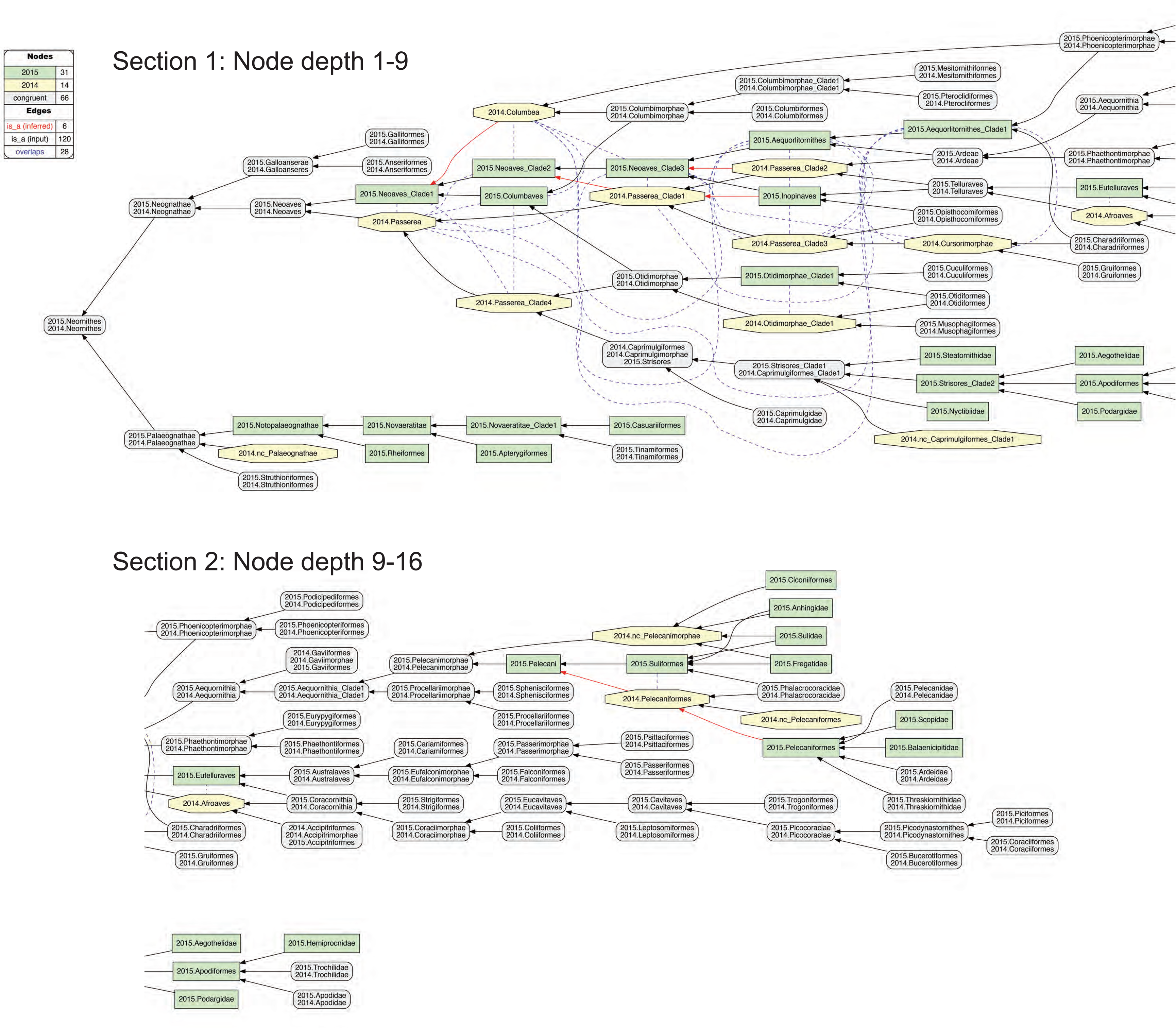
Alignment visualization for the 2015./2014.Neornithes alignment, under whole-concept resolution, ranging from the root to the ordinal level (with exceptions where needed), and with 28 overlapping relationships. Compare with Figs. 7, 9, and 10; see text for further detail. The reasoner infers 8,051 logically implied articulations that constitute the set of MIR. See also S16 File set.

**Fig 12.**
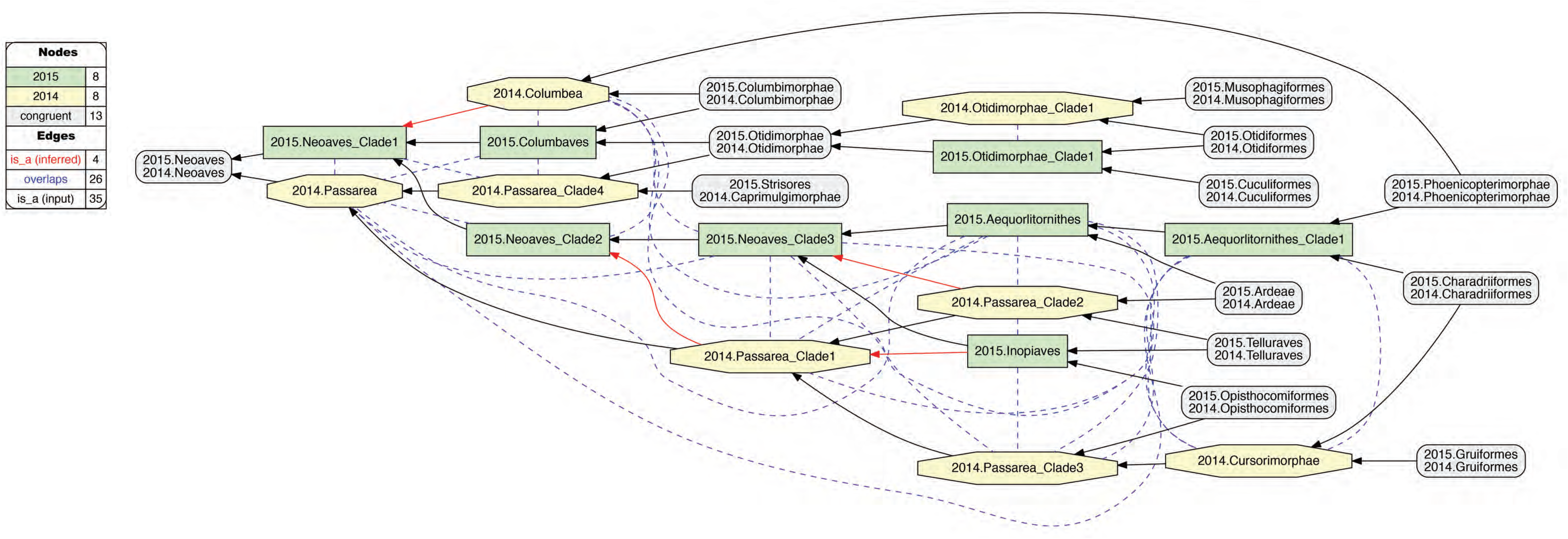
Alignment visualization for the 2015./2014.Neoaves alignment, under whole-concept resolution, limited tothe main conflict region, and with 26 overlapping relationships. Compare with Fig. 11. The reasoner infers 441 logically implied articulations that constitute the set of MIR. See also S17 File set.

**Fig 13.**
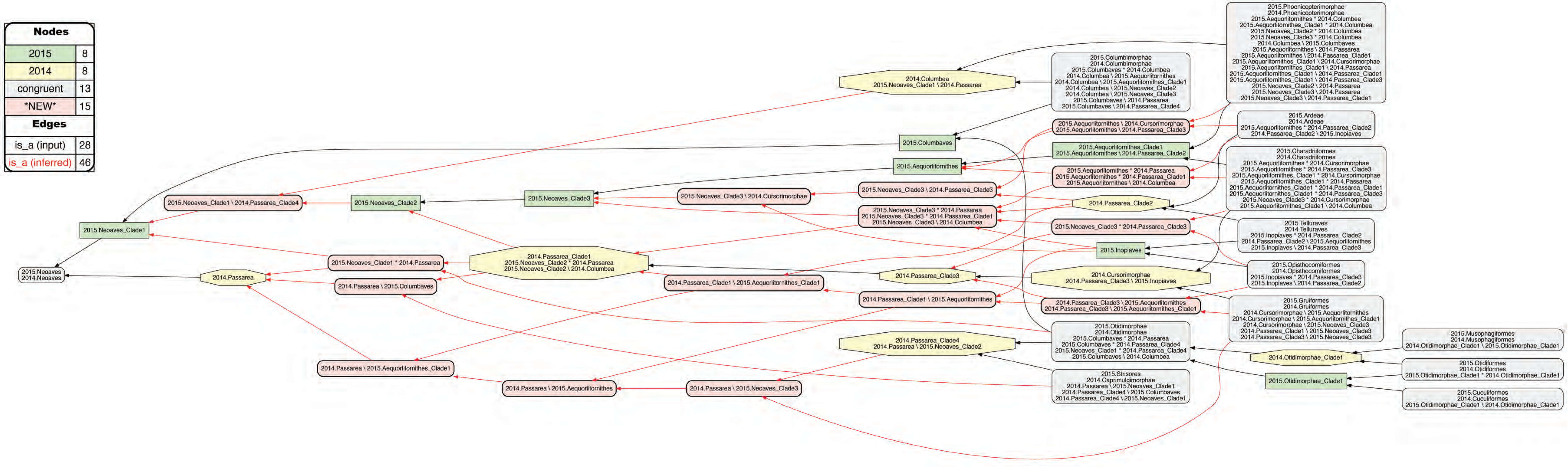
Alignment visualization for the 2015./2014.Neoaves alignment, under split-concept resolution, limited to the main conflict region, and resolving the 26 overlapping relationships. Compare with Fig. 12; the 15 salmon-colored regions are *only* identifiable via split-concept resolution labels. Compare with Fig. 12 and Table 4. See text for further detail. The reasoner infers 441 logically implied articulations that constitute the set of MIR. See also S18 File set.

The two author teams sampled four family-level concepts congruently for the 2015./2014.Pelecanimorphae alignment region (Fig. 7). However, 2015.PEA’s phylogeny entails six additional family-level concepts that have no apparent match in 2014.JEA. Moreover, the latter authors recognize only one rather inclusive order-level concept, 2014.Pelecaniformes, under which all four family-level concepts are subsumed, including 2014.Phalacrocoracidae. In contrast, 2015.PEA an intensionally less inclusive – though more comprehensively sampled – concept of 2015.Pelecaniformes, and place their congruent 2015.Phalacrocoracidae in the order-level concept 2015.Suliformes. This represents the first instance of plausibly rejecting the proposition: “Had 2014. JEA sampled 2014.Phalacrocoracidae, they would have assigned this concept to 2014.Suliformes”. The assertion is no longer counter-factual: 2014.JEA *did* sample the corresponding child concept (2014.Phalacrocoracidae), but did *not* assign it to a parent concept separate from 2014.Pelecaniformes. Accordingly, we obtain three overlapping, ‘cascading’ articulations between concepts that form the 2015.Suliformes higher-level topology and 2014. Pelecaniformes. Meanwhile, the uniquely sampled 2015.Ciconiiformes are subsumed under 2014. Pelecanimorphae which has relaxed parent coverage.

Within 2015.Pelecaniformes, we obtain five additional overlapping articulations between five concepts that make up the 2015/2014 supra-familial topologies in this alignment (Fig. 7). Here the conflict is due to the differential assignment of 2015./2014.Pelecanidae. Specifically, 2015.PEA inferred a sister relationship of 2015.Pelecanidae with 2015.Balaenicipitidae, for which 2014.JEA have no sampled match. Meanwhile, the latter authors inferred a sister relationship of 2014. Pelecanidae with 2014.Ardeidae. The latter concept *is* matched in 2015.PEA with 2015. Ardeidae, though not as the most immediate sister concept of 2015.Pelecanidae. Of course, we may posit that a 2015.Ardeidae/2015.Pelecanidae sister relationship is what 2015.PEA *would* have obtained, had these authors not also sampled 2015.Balaenicipitidae and 2015.Scopidae. But they did, and hence obtained two clade concepts that include 2015.Pelecanidae yet exclude 2015.Ardeidae; i.e., 2015.Pelecanoidea_Clade1 and 2015.Pelecanoidea_Clade2. While relaxing parent coverage for 2014.Pelecaniformes_Clade2 could serve to mitigate this conflict, we deem the overlapping relationship to better represent 2015.PEA’s phylogenomic signal, which happens to ‘break up’ the lowest supra-familiar clade concept supported by 2014.PEA.

The 2015.Passeri/2014.Passeriformes_Clade2 alignment is another instance where relaxing parent coverage can only partially mitigate conflict (Fig. 8). In this case, 2015.PEA and 2014.JEA sampled two sets of family-level concepts that are wholly exclusive of each other, *except* for 2015./2014.Corvidae. Regarding the only two additional family-level concepts recognized in 2014. JEA – i.e., 2014.Passeridae and 2014.Thraupidae – we may posit counter-factually that these would be subsumed under 2015.Passeroidea with relaxed coverage [47]. However, further assertions of congruence are difficult to justify, given the limited sampling of 2014.JEA. Thus, in our current representation, 2014.Passeriformes_Clade2 shows an overlapping relationship with 2015. Passeroidea, its immediate parent 2015.Passerida, and also with 2015.Corvoidea.

A single yet significant instance of overlap occurs just within the congruent parent concepts 2015./2014.Telluraves (Fig. 9). Two levels below this paired parent region, both author teams recognize three congruent children; viz. 2015./2014.{Australaves, Coracornithia, Accipitrimorphae/Accipitriformes}. However, 2015.Prum group the former two concepts under 2015.Eutelluraves, with 2015.Accipitriformes as sister; whereas 2014.JEA cluster the latter two concepts under 2014.Afroaves, with 2014.Australaves as sister. This amounts to the first occurrence of conflict that *cannot* justifiably be resolved by relaxing parent coverage, but instead reflects divergent phylogenomic signals.

### Whole-concept and split-concept resolution

How to *speak* of such overlap? In Fig. 9, as in the preceding Figs. 7 and 8 that showed overlapping relationships, we only utilize clade concept labels that are consistent with each input phylogeny. In the resulting alignment, the articulation 2015.Eutelluraves >< 2014.Afroaves is visualized as a dashed blue line between these regions that retain the same extension through the input-reasoning-output transition. Yet Fig. 9 also specifies the extent of regional overlap at the next lower level. Accordingly, the paired concept region 2015./2014.Coracornithia is that which is actually subsumed under each of the overlapping parents – as indicated by the two inclusion arrows that extend ‘upward’ from this region. The other two paired child regions are respectively members of one parent region only.

If we call the input regions 2015.Eutelluraves “A” and 2014.Afroaves “B” in this example, we can use the following syntax to identify output regions that result from overlapping input concepts [26]: A*B (read: “A *and* B”) constitutes the output region shared by two parents, whereas A\b (“A, *not* b”) and B\a (“B, *not* a”) are output regions with only one parent. We call this more granular syntax *split-concept resolution* (“merge concepts” in [26]), as opposed to *whole-concept resolution* which preserves the syntax and granularity provided by the input concept labels.

In Fig. 10, the 2015./2014.Telluraves overlap is represented with split-concept resolution. This eliminates the need to visualize a dashed blue line between 2015.Eutelluraves and 2014.Afroaves (Fig. 9). Moreover, in this case the split-concept resolution syntax is redundant or unnecessary, because each of the three resolved regions under “A” (2015.Eutelluraves) and “B” (2014.Afroaves) is congruent with two regions already labeled in the corresponding input phylogenies. We will see, however, that this granular syntax is essential for verbalizing the outcomes of more complex alignments that contain many overlapping regions.

### Zooming in on the neoavian explosion

The remaining 26 instances of overlap are shown under different alignment visualizations in Figs. 11–13. They occur 1-5 levels below the congruent concept pair 2015./2014.Neoaves, and jointly define the primary region of phylogenomic conflict between 2015.PEA and 2014.JEA that presumably inspired the term “neoavian explosion”. Because parent coverage was selectively applied at lower levels, none of the 26 overlaps in the alignment are caused by differential child sampling. Instead, they represent genuine phylogenomic conflict in the higher-level arrangement of congruent sets of children.

Our Fig. 11 is intended to be an RCC–5 alignment *analogue* to figure 1 in [3] – reproduced here with permission as Fig. 14. The alignment generally reaches from the root to the ordinal level, and to the family level in the two subregions where order-level concepts are conflicting (see Fig. 4 and S12 File set). The visualization offers an intuitive signal of the abundance and distribution of in/congruence throughout the alignment. In all, 66/111 regions (59.5%) are congruent, of which 22 are located in the 2015./2014.Telluraves (though see Figs. 9 and 10); 15 are contained in the 2015./2014.Ardeae (including four family-level regions); and 5 are part of the 2015./2014.Columbimorphae. Outside of the 2015./2014.Neoaves, 8 such regions are present. In other words, the two phylogenies are congruent at the highest level and also in several intermediate regions above the ordinal level.

**Fig 14.**
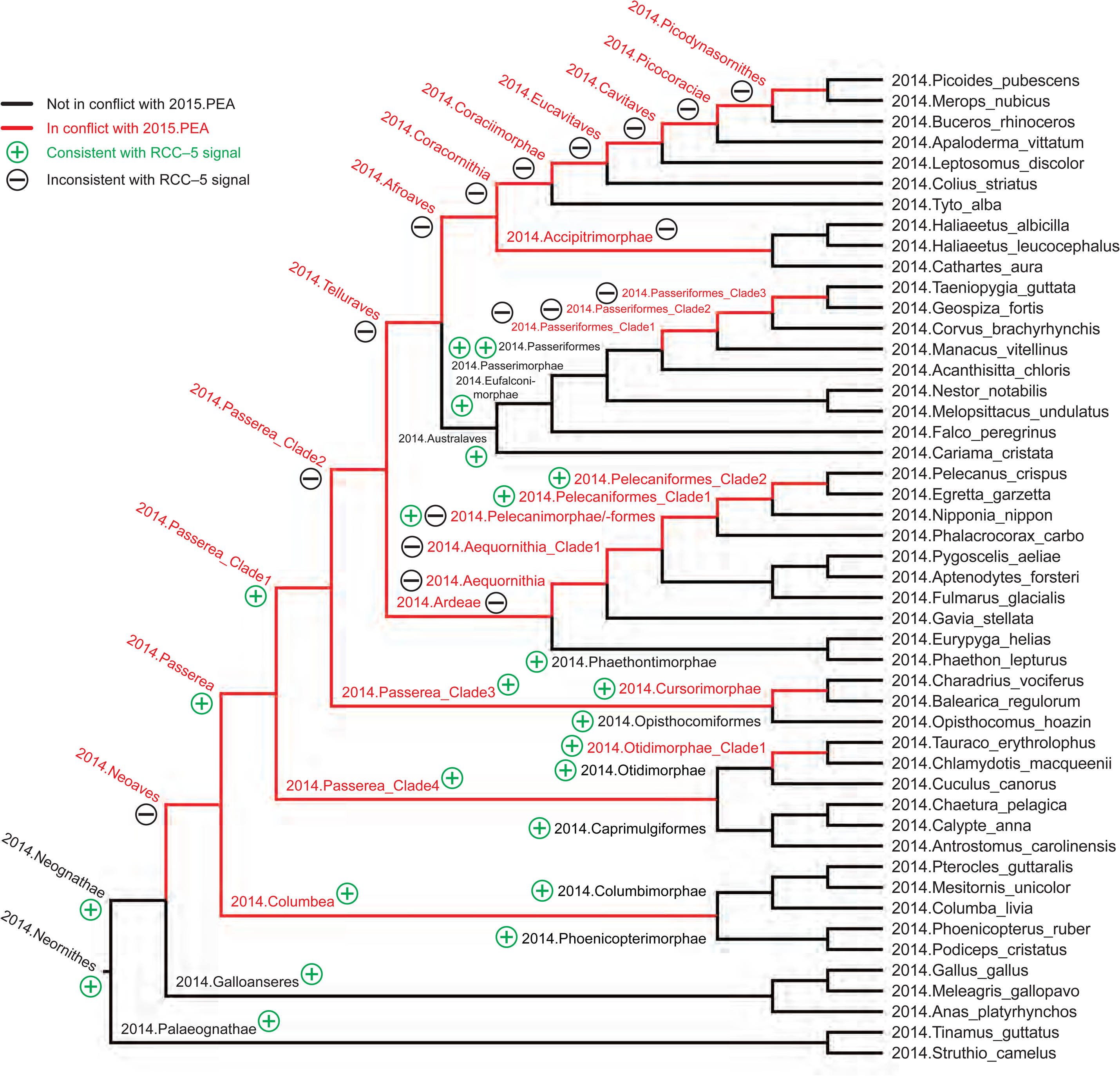
Conflict visualization for Avian phylogenomic relationships, using the method of [11, 15, 23], with 2014.JEA as the primary source phylogeny and 2015.PEA as the alternative. Black edges indicate concordance, whereas red edges signal conflict. Clade and terminal concept labels are added in accordance with the present study. Moreover, consistency or inconsistency of the edge concordance/conflict analysis with the RCC–5 alignments (Figs. 7 to 13) are signaled via a green “+” circle and a black “−” circle, respectively. See also S19 File.

Figure 12 shows just the neoavian explosion region under whole-concept resolution. Each phylogeny contributes 21 input concepts to this ‘zoomed-in’ alignment, which yields 13 congruent regions. Of these, only 2015./2014.Neoaves and 2015./2014.Otidimorphae represent non-terminal concepts.

Unpacking the complexity of this conflict region requires a stepwise analysis. From the perspective of 2015.PEA, the 2015.Neoaves are split into a sequence of three unnamed, higher-level clade concepts, i.e. 2015.{Neoaves_Clade1, Neoaves_Clade2, Neoaves_Clade3}, with 2015.{Strisores, Columbaves, Gruiformes} as corresponding sister concepts. The two children of 2015.Neoaves_Clade3 are 2015.{Aequorlitornithes, Inopinaves}. The authors accept the nomenclature of [44] for 2015.Strisores, with is congruent with 2014.Caprimulgimorphae and the 2015. /2014.Gruiformes as congruent as well. However, the remaining six high-level concepts of 2015. PEA are exceedingly poorly aligned with the two highest-level neoavian concepts of 2014. JEA, i.e. 2014.{Columbea, Passerea}, and also with any of the four unnamed clade concepts below 2014.Passerea. In particular, consecutive node sequence 2015.{Neoaves_Clade3, Aequorlithornites, Aequorlithornites_Clade1} participates in 16/26 overlaps, as summarized in Table 3. Loosely corresponding to this sequence are the concepts 2014.{Passerea_Clade1, Passerea_Clade2, Cursorimorphae}, jointly with 10 overlaps. As shown in Fig. 12, these overlaps are grounded in the incongruent assignment of five paired, lower-level concept regions; viz. 2015. /2014.{Ardeae, Charadriiformes, Opisthocomiformes, Phoenicopterimorphae, Telluraves}. Two strongly conflicting placements contribute most to the number of overlaps: (1) 2015. /2014.Charadriiformes in 2015.Aequorlithornites_Clade1 (sister to 2015. Phoenicopterimorphae) versus 2014.Cursorimorphae (sister to 2014.Gruiformes); and (2) 2015. /2014.Phoenicopterimorphae in 2015.Aequorlithornites_Clade1 versus 2014.Columbea (sister to 2014.Columbimorphae). Indeed, if one had to single out one concept inferred in 2015. PEA that causes the most topological incongruence with 2014.JEA - this would indicate the newly proposed yet unnamed 2015. Aequorlithornites_Clade1, consisting of certain “waterbirds”. This 2015.PEA concept, together with its four superseding parents – of which only one is explicitly named – ‘triggers’ 20/26 overlaps with the phylogenomic tree of 2014.JEA.

**Table 3.** Overview of 26 pairwise 2015/2014 concept overlaps in main neoavian conflict region. See also Figs. 11 and 12.

| Clade concept label | 2014.<br>Passerea | 2014.<br>Columbea | 2014.Passerea<br>_Clade3 | 2014.Cursori-<br>morphae | 2014.Passerea<br>_Clade1 | 2014.Passerea<br>_Clade2 | 2014.Passerea<br>_Clade4 | 2014.Otidimor-<br>phae_Clade1 | Totals |
| --- | --- | --- | --- | --- | --- | --- | --- | --- | --- |
| 2015.Aequorlithornites | >< | >< | >< | >< | >< | >< |  |  | 6 |
| 2015.Aequorlithornites_Clade1 | >< | >< | >< | >< | >< |  |  |  | 5 |
| 2015.Neoaves_Clade3 | >< | >< | >< | >< | >< |  |  |  | 5 |
| 2015.Columbaves | >< | >< |  |  |  |  | >< |  | 3 |
| 2015.Neoaves_Clade1 | >< |  |  |  |  |  | >< |  | 2 |
| 2015.Neoaves_Clade2 | >< | >< |  |  |  |  |  |  | 2 |
| 2015.Inopinaves |  |  | >< |  |  | >< |  |  | 2 |
| 2015.Otidimorphae_Clade1 |  |  |  |  |  |  |  | >< | 1 |
| Totals | 6 | 5 | 4 | 3 | 3 | 2 | 2 | 1 | 26 |

Two additional ‘centers’ of conflict are identifiable in Fig. 12. The first, embedded in the aforementioned region, concerns the alignment of the two concepts 2015.Inopinaves and 2014. Passerea_Clade2, which share the child regions 2015./2014.Telluraves, yet differentially accommodate the congruent regions 2015./2014.Ardeae and 2015./2014.Opisthocomiformes. This relationship further contributes to the ‘cloud’ of overlaps along the respective 2015. Neoaves_Clade{1-3}/Aequorlithornites/_Clade1 and 2014.Passerea/_Clade{1- 3}/Cursorimorphae concept topology chains. Second, the two paired regions 2015.2014.Columbimorphae and 2015./2014.Otidimorphae are incongruently assigned to three overlapping parents, i.e. 2015.Columbaves and 2014.{Columbea, Passerea_Clade4}. From the perspective of 2015.PEA, then, 2014.JEA's highest-level neoavian bifurcation of 2014.Columbea and 2014.Passerea is arguably the most salient subregion of conflict, as these two concepts alone participate in 11 overlaps. A third, relatively minor incongruence concerns the relative placement of three paired ordinal concept regions within the 2015./2014Oditimorphae.

### Split-concept resolution for the neoavian explosion

In Fig. 13, the same ‘zoomed-in’ alignment is shown under split-concept resolution. The more granular set of clade concept labels permits identifying all output regions created by the 26 overlaps of the neoavian explosion; see also Table 4. The entire set consists of 78 labels; i.e., 26 labels for each split-resolution product {A*B, A\b, B\a} corresponding to one input region overlap. Interestingly, and unlike the outcome displayed in Fig. 10, not all of these split-concept resolution labels are semantically redundant with those provided in the input, although most are. Specifically, 51 labels are generated ‘in addition’ for the 12 terminal congruent regions (compare with Fig. 12). These are indeed unnecessary, in the sense that they are semantic synonyms for regions already appropriately identified in the input. However, the relative *number* of additional labels generated per input-identified region is telling. This number will be highest for those (terminal) regions whose differential placements into higher-level regions are the primary drivers of incongruence in the alignment. As analyzed above, these are: 2015./2014.{Phoenicopterimorphae, Charadriiformes, Columbimorphae}, respectively with 14, 8, and 7 additional labels. Six redundant split-concept resolution labels are further produced for input regions that are unique to one phylogeny; e.g., 2014.Columbea is understandably also labeled 2015.Neoaves_Clade1 \ 2014. Passerea (where the “\” means: not).

**Table 4.** Overview of 15 newly inferred split-concept resolution regions and labels (or label clusters) for the neoavian conflict region that lack appropriate input clade concept labels. See also Fig. 13. For each split-concept resolution label (or label cluster), we provide the two immediate children or constituent concepts 1 and 2 – i.e., what is jointly subsumed ‘underneath’ the split – as well as the set of lower-level concept regions (using whole-concept resolution labels) that are differentially distributed by the split between the two source phylogenies. * = Two children listed.

| # | Split-concept label(s) | Constituent clade concept 1 | Constituent clade concept 2 | Lower-level concept regions differently assigned by the split |
| --- | --- | --- | --- | --- |
| 1 | 2015.Neoaves_Clade1 * 2014.Passarea_Clade4 | 2015.Neoaves_Clade2 | 2014.Columbea | 2015./2014.Otidimorphae, 2015.Strisores/2014.Caprimulgimorphae |
| 2 | 2015.Neoaves_Clade1 * 2014.Passarea | 2015./2014.Otidimorphae | 2014.Passarea_Clade1 | 2015.Strisores/2014.Caprimulgimorphae, 2014.Columbea |
| 3 | 2014.Passarea \ 2015.Columbaves | 2015.Strisores/2014.Caprimulgimorphae | 2014.Passarea_Clade1 | 2015./2014.Columbimorphae, 2015./2014.Otidimorphae |
| 4 | 2014.Passarea \ 2015.Aequorlornithes_Clade1 | 2015.Aequorlornithes \ 2014.Passarea | 2014.Passarea_Clade1 \ 2015.Aequorlornithes_Clade1 | 2015./2014.Charadriiformes, 2015./2014.Phoenicopterimorphae |
| 5 | 2014.Passarea \ 2015.Aequorlornithes | 2014.Passarea \ 2015.Neoaves_Clade3 | 2014.Passarea_Clade1 \ 2015.Aequorlornithes | 2015./2014.Ardeae, 2015./2014.Charadriiformes, 2015./2014.Phoenicopterimorphae |
| 6 | 2015.Neoaves_Clade3 \ 2014.Cursorimorphae | 2015.Inopinaves | 2015.Neoaves_Clade3 \ 2014.Passarea_Clade3 | 2015./2014.Charadriiformes, 2015./2014.Gruiformes |
| 7 | 2014.Passarea \ 2015.Neoaves_Clade3 | 2015./2014.Gruiformes | 2014.Passarea_Clade4 | 2015./2014.Ardeae, 2015./2014.Charadriiformes, 2015./2014.Opisthocomiformes, 2015./2014.Phoenicopterimorphae, 2015./2014.Telluraves |
| 8 | 2014.Passarea_Clade1 \ 2015.Aequorlornithes_Clade1 | 2014.Passarea_Clade1 \ 2015. Aequorlornithes | 2014.Passarea_Clade2 | 2015./2014.Charadriiformes, 2015./2014.Phoenicopterimorphae |
| 9 | {2015.Neoaves_Clade3 \ 2014.Columbea, 2015.Neoaves_Clade3 * 2014.Passarea, 2015.Neoaves_Clade3 * 2014.Passarea_Clade1} | {2015.Aequorlornithes \ 2014.Columbea, 2015.Aequorlornithes * 2014.Passarea, 2015.Aequorlornithes * 2014.Passarea_Clade1}, 2015.Neoaves_Clade3 * 2014.Passarea_Clade3 * | 2015.Inopinaves, 2014.Passarea_Clade2 * | 2015./2014.Columbimorphae, 2015./2014.Gruiformes, 2015./2014.Otidimorphae, 2015./2014.Phoenicopterimorphae, 2015.Strisores/2014.Caprimulgimorphae |
| 10 | 2015.Neoaves_Clade3 \ 2014.Passarea_Clade3 | {2015.Aequorlornithes \ 2014.Cursorimorphae, 2015.Aequorlornithes \ 2014.Passarea_Clade3} | 2014.Passarea_Clade2 | 2015./2014.Ardeae, 2015./2014.Charadriiformes, 2015./2014.Gruiformes, 2015./2014.Opisthocomiformes, 2015./2014.Phoenicopterimorphae, 2015./2014.Telluraves |
| 11 | 2014.Passarea_Clade1 \ 2015.Aequorlornithes | 2014.Passarea_Clade3 \ 2015.Aequorlornithes, 2014.Passarea_Clade3 \ 2015.Aequorlornithes_Clade1} | 2015.Inopinaves | 2015./2014.Ardeae, 2015./2014.Charadriiformes, 2015./2014.Gruiformes, 2015./2014.Opisthocomiformes, 2015./2014.Phoenicopterimorphae, 2015./2014.Telluraves |
| 12 | {2015.Aequorlornithes \ 2014.Cursorimorphae, 2015.Aequorlornithes \ 2014.Passarea_Clade3} | 2015./2014.Phoenicopterimorphae | 2015./2014.Ardeae | 2015./2014.Gruiformes, 2015./2014.Opisthocomiformes |
| 13 | {2015.Aequorlornithes \ 2014.Columbea, 2015.Aequorlornithes * 2014.Passarea, 2015.Aequorlornithes * 2014.Passarea_Clade1} | 2015./2014.Ardeae | 2015./2014.Charadriiformes | 2015./2014.Columbimorphae, 2015./2014.Gruiformes, 2015./2014.Opisthocomiformes, 2015./2014.Otidimorphae, 2015./2014.Phoenicopterimorphae, 2015.Strisores/2014.Caprimulgimorphae, 2015./2014.Telluraves |
| 14 | 2015.Neoaves_Clade3 * 2014.Passarea_Clade3 | 2015./2014.Charadriiformes | 2015./2014.Opisthocomiformes | 2015./2014.Ardeae, 2015./2014.Gruiformes, 2015./2014.Phoenicopterimorphae, 2015./2014.Telluraves |
| 15 | {2014.Passarea_Clade3 \ 2015.Aequorlornithes, 2014.Passarea_Clade3 \ 2015.Aequorlornithes_Clade1} | 2015./2014.Gruiformes | 2015./2014.Opisthocomiformes | 2015./2014.Ardeae, 2015./2014.Charadriiformes, 2015./2014.Phoenicopterimorphae |

In contrast, the remaining 21 split-concept resolution labels identify 15 salmon-colored alignment regions – 11 uniquely and 4 redundantly with 2–3 labels each – for which there are *no* suitable labels in either of the phylogenomic input trees (Table 4). Forty-six additional articulations are inferred to align these regions to those displayed in Fig. 12, under whole-concept resolution. To reiterate: these novel regions are not congruent with any clade concepts recognized by the source phylogenies. However, they are needed if we wish to express how exactly the authors' respective clade concepts overlap.

Consider the example of the higher-level split-concept resolution region 2015.Neoaves_Clade1 * 2014.Passerea (A*B). This region has two obvious parents – i.e., 2015.Neoaves_Clade1 and 2014. Passerea – and entails just those children that each shares: 2015./2014Otidimorphae and 2014. Passerea_Clade1. Excluded from this intersection are: 2015.Neoaves_Clade \ 2014.Passerea (A\b) in the view of 2015.PEA, and 2014.Passerea \ 2015.Neoaves_Clade1 (B\a) in the view of 2014. JEA. The A\b split is also labeled 2014.Columbea, whereas the B\a split can be labeled 2015.Strisores or 2014.Caprimulgimorphae. Table 4 specifies this kind of partitioning and labeling for each of the 15 newly inferred split-concept resolution regions.

Three distinct reference services are gained by generating the split-concept resolution labels. First, in cases where no whole-concept resolution labels are available, we obtain appropriately short and consistent labels to identify the split regions caused by overlapping clade concepts. Of course, we could also label 2015.Neoaves_Clade1 * 2014.Passerea (A*B) as “2015./2014.Otidimorphae and 2014.Passerea_Clade1”. That is, we could provide a list of all whole-concept resolution children subsumed under split-concept resolution region. But this approach will frequently lead to long lists of labels if the phylogenies are large and splits are topologically complex. Second, the {A*B, A\b, B\a} triplets have an explanatory, provenance-signaling function, by using the same syntactic set of input labels (A, B) to divide topologically complementary alignment regions of an overlap. If we focus on one label of a triplet, we can find the two complements and thereby systematically explore the 'reach' of each split in the alignment. Third, the clade concept labels (A, B) used in the split-concept resolution labels will be exactly those that identify incongruent, overlapping regions across the source phylogenies. In other words, there is a special cognitive value in using this convention.

### Analysis of clade *name* performance

We can now also ask, specifically with regards to the alignment shown in Fig. 11, to what extent the clade names (syntax) used by the two author teams succeed or fail to identify congruent and incongruent concept regions (semantics). Such name:meaning (read: “name-to-meaning”) analyses were carried out in three previous alignment use cases, with rather unfavorable outcomes for the respective names in use [14, 32, 51]. Here, the 97 x 83 input concepts yield a set of 8,051 MIR (S16D File). Of these, 384 MIR involve one of four “no coverage” regions added to 2014.JEA concepts at this level. Because the latter are in effect placeholders for relaxing a constraint, we restrict the name:meaning analysis to the remaining 7,667 MIR (Table 5).

**Table 5.** Clade name-to-RCC–5 relationship reliability analysis for the higher-level neoavian explosion alignment. Relationship data are derived from the set of MIR corresponding to Fig. 11 and the S16D File.

| Relationship (RCC-5 / clade name) | == | > | < | >< | ! | Totals |
| --- | --- | --- | --- | --- | --- | --- |
| Same clade name | 62 | 0 | 1 | 1 | 0 | 64 |
| Different clade names | 7 | 625 | 691 | 27 | 6,253 | 7,603 |
| Totals | 69 | 625 | 692 | 28 | 6,253 | 7,667 |

Interestingly, the clades names used by the respective author teams fare rather well in comparison, because only nine of 7,667 pairings in the MIR (0.12%) are unreliable as identifiers of in-/congruence of the respective RCC–5 articulation. In seven instances, two congruent concepts have different names. Four of these merely involve changes in name endings, such as 2015. Accipitriformes == 2014.Accipitrimorphae, 2015.Galloanserae == 2014.Galloanseres, or 2015. Pteroclidiformes == 2014.Pterocliformes. The articulation 2015.Pelecaniformes < 2014. Pelecaniformes is the single instance in which the meaning of the same name is less inclusive in one source (Fig. 7). Lastly, the relationship 2015.0tidimorphae_Clade1 >< 2014.0tidimorphae_Clade1 involves the same name in an overlapping articulation (Figs. 12 and 13), though we have to recognize that neither author team actually provided this clade name (see Methods).

In summary, the clade concept names used respectively by 2015.PEA and 2014.JEA very rarely provide an *incorrect* signal regarding semantic in-/congruence. This desirable outcome seems to reflect on each author team’s recognition that their newly inferred clade concepts within the 2015. /2014.Neoaves merit appropriately unique names. However, not providing misleading names is not enough if we intend to speak of the various kinds of conflict that exist between the different phylogenomic signals.

### Comparison with other conflict visualizations

We now compare these results with conflict analysis and visualization tools created for the Open Tree of Life project (OToL) – a community-curated platform for synthesizing our knowledge of phylogeny [13, 22, 23, 24]. The approach we employ is explained and applied in [11, 15, 23, 52, 53]. The method starts off with ‘normalizing’ all terminal names in the source trees to a common taxonomy [24]. Having the same normalized terminal name means taxonomic concept congruence (==) at that level. To assess conflict from the perspective of one rooted input tree (A), a source edge *j* of that tree is taken to define a rooted bipartition *S*(*j*) = *S*_in_ | *S*_out_, where *S*_in_ and *S*_out_ are the tip sets of the ingroup and outgroup, respectively. The algorithm progresses sectionally from the leaves to the root. Concordance or conflict for a given edge *j* in tree A with that of tree B is a function of the relative overlap of the corresponding tip sets, as follows [23]. Concordance between two edges in the input trees A and B is obtained when B_in_ is a proper subset (⊂) of A_in_ *and* B_out_ ⊂ A_out_. On the other hand, two edges in trees A and B are conflicting if *none* of these sets are empty: A_in_ intersects (∩) with B_in_, A_in_ ∩ B_out_, or B_in_ ∩ A_out_. In other words, conflict means that there is reciprocal overlap in the ingroup and outgroup bipartitions across the two trees.

We applied this approach in both directions, i.e. starting with 2014.JEA as primary source and identifying edges therein that conflict with those of 2015.PEA, and vice-versa. The visualizations are shown in Figs. 14 and 15, respectively. Our examination of these results remains qualitative and in service of the main theme.

**Fig 15.**
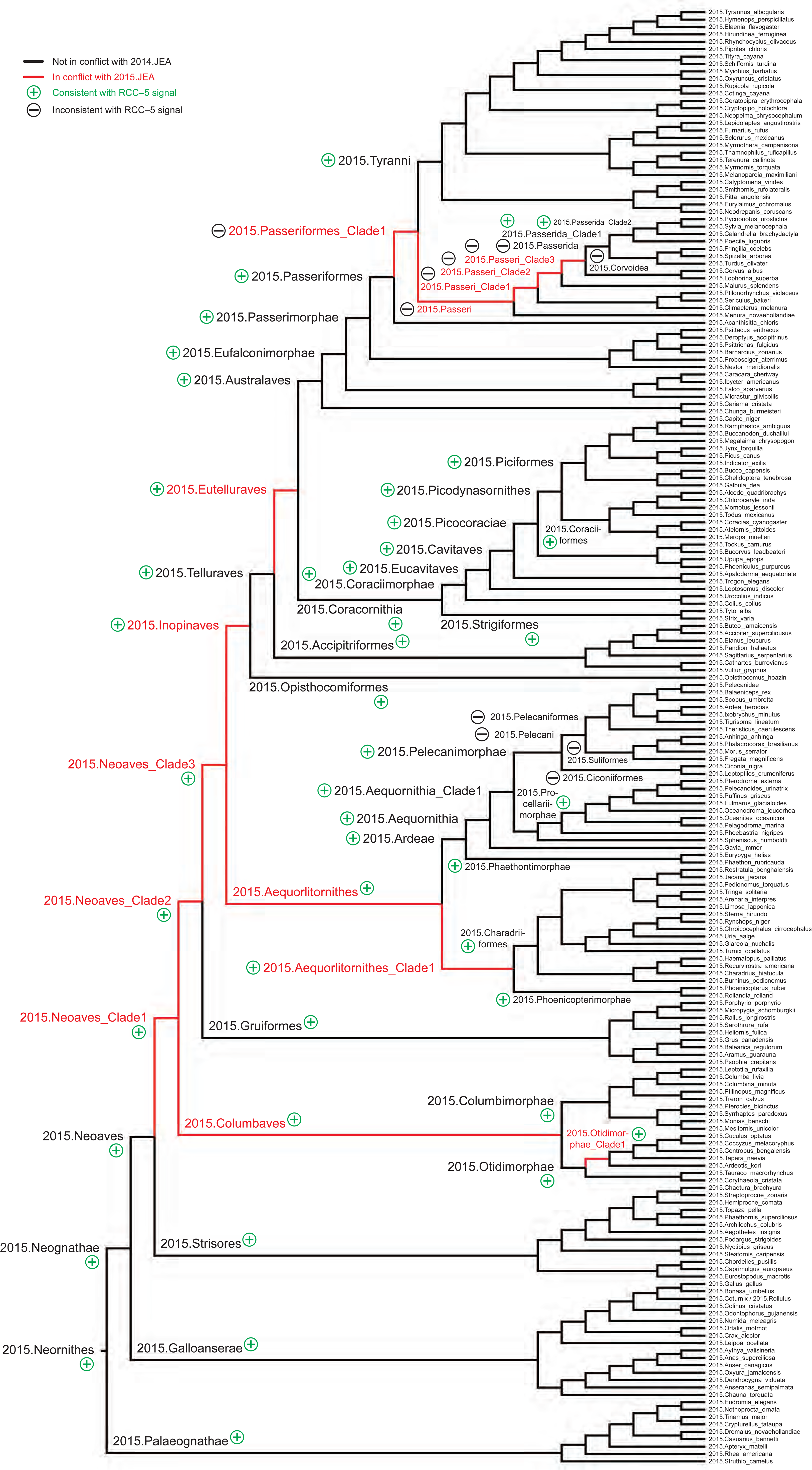
Conflict visualization for Avian phylogenomic relationships, using the method of [11, 15, 23], with 2015.PEA as the primary source phylogeny and 2014.JEA as the alternative. Display conventions as in Fig. 14. See also S20 File.

Most of the red edges particularly in Fig. 15 – which is based on the nearly four times more densely sampled tree sec. 2015.PEA – are consistent with the overlapping RCC–5 relationships shown in Figs. 7 to 13. However, within the 2015.Pelicanimorphae, certain RCC–5 overlaps (Fig. 7) are not recovered (“false positives”), whereas within the 2015.Passeriformes, numerous edges are shown as conflicting (“false negatives”) that are merely congruent refinements according to the RCC–5 alignment (Fig. 8).

Using this method with the less densely sampled tree sec. 2014.JEA as the base topology creates certain inconsistencies that are instructive (Fig. 14). Here, a much larger subset of the topology ‘backbone’ – starting with the 2014.Neoaves, and continuing via 2014.Passerea, 2014.Telluraves, 2014. Afroaves, to (e.g.) 2014.Picodynasornithes – is inferred by the algorithm as conflicting. Many of these red edges lack reciprocal correspondence with the visualization grounded in 2015. JEA’s phylogeny. For instance, 2014.{Neoaves, Ardeae, Coracornithia} are shown as conflicting edges in Fig. 14, when 2015.{Neoaves, Ardeae, Coracornithia} are concordant edges in Fig. 15. These inconsistencies are caused by the addition of terminals sec. 2015.PEA that have no suitably granular matches in 2014.JEA’s sampled tips and tree, which means that they will attach as children to a higher-level parent represented in the OToL taxonomy. The latter is assembled separately [24], and is used to place terminals that are differentially sampled between to sources. For instance, 2015.Ciconiiformes – which has no close match in 2014.JEA - may end up attaching as a child of 2014.Neognathae instead of 2014.Pelecanimorphae (Fig. 7). The taxonomy in effect assumes the responsibility of representing concept intensionality, in lieu of trained expert judgment. However, at the time of analysis, the common OToL taxonomy lacked a name/concept for “Neoaves”. This means that the 2015./2014.Neoaves ingroup/outgroup bipartitions will conflict in the placement of 2015.Ciconiiformes, and hence show this conflict in Fig. 14 but not in Fig. 15.

## Discussion

### Key phylogenomic conflict representation conventions

We first review the key conventions of our RCC–5 multi-phylogeny alignment approach, before discussing the services that can be derived from the alignments.

1. Using the taxonomic concept label convention of [14] allows us to individuate each concept entailed in 2014.JEA and 2015.PEA, even if the taxonomic or clade concept *name* components are identical, as in 2015.Pelicaniformes < 2014.Pelicaniformes.
2. Because our main intention is to represent phylogenomic congruence and conflict across these inferred phylogenies, there is no need to speak of sameness in any profound sense, such as referring to the “same {clades, nodes, species, taxa}”. Such language is best used once we shift from (1) logically modeling similarities and differences between human-made phylogenomic theories to (2) hopefully but not necessarily robust evolutionary inferences. Speaking exclusively of *concepts* and their relative congruence in our RCC–5 alignments is appropriate because we thereby do not blur the lines between two important communication goals – i.e., systematic conflict representation and evolutionary inference – that are each best met by maintaining complementary manners of speaking, as much as possible [21].
3. Linking concepts via *is_a* (parent/child) relationships permits the assembly of single-source hierarchies, whereas the RCC–5 articulations express the relative congruence of concept regions across multi-source hierarchies. Uncertainty can be accommodated via disjunctions of the base five relations, i.e., by asserting subsets of the R_32_ relationship lattice [33].
4. Under the default logic reasoning constraint of parent coverage, differential child-level sampling will propagate up to yield incongruent relationships among the respective parent-level clade concepts [14, 26, 29]. Local relaxation of the coverage constraint can mitigate this effect. However, this requires an explicit application of trained expert judgment [30], reflected in assertions of input articulations that, in effect, stipulate counter-factual circumstances. We can thereby indirectly model *intensional* (property-based) node concept definitions in the RCC–5 alignments, and hence obtain select instances of higher-level clade concept congruence in spite of incongruent low-level sampling (Figs. 1–4).
5. Because every clade concept region to be aligned requires at least a functional label – with a syntax suitable for human communication – we need to supply such labels for clade concepts which the original sources do not explicitly identify by name. If no published name is applicable, a pragmatic solution is to utilize the next available higher-level name and add the suffix “_Clade#”, as in 2015.Neoaves_Clade1 or 2014.Passerea_Clade3 (Tables 1 and 2).
6. In some instances, an original source may provide a clade concept label, such as 2015. Eurylaimidae, that corresponds only to a non-monophyletic region of the source’s phylogenomic topology. Representing such label-to-clade-concept mismatches logically amounts to providing yet another alignment between (1) the reference classification from which the label was adopted and (2) the phylogeny to which it is incongruently applied (Figs. 5 and 6).
7. Partitioning of the global alignment may be required to manage the use case complexity, given the current logic reasoner performance. Consistent alignments of higher-level concept hierarchies, as shown in Fig. 11, can be derived from this bottom-up approach [14, 28, 51].
8. Overlapping relationships among higher-level clade concepts can be represented using either whole-concept or split concept resolution (compare Figs. 9 and 10; Figs. 12 and 13). The latter option provides a uniquely granular and informative syntax to partition and label the alignment regions created by phylogenomic concept overlap (Tables 3 and 4).
9. The reasoner-inferred MIR allow us to quantify all pairwise instances where the *names* used by each source succeed, or fail, in matching the semantic signal of the RCC–5 relationships and concept labels (Table 5).
10. The outcomes of this approach can be compared with other conflict representation methods, such as those created for the OToL project [11, 15, 23]. This is particularly illustrative in cases where differential sampling of low-level concepts generates unequal assessments between the OToL and RCC–5 approaches regarding higher-level clade concept identity and concordance (Figs. 14 and 15).

### Verbal and visual knowledge representation services

What can we gain from this approach, both narrowly for the neoavian explosion use case and more generally for the future data integration in systematics?

Data representation designs have inherent trade-offs. Unlike other semi-/automated phylogenomic conflict visualization methods [13, 23, 24], using the above approach requires extensive *upfront* application of human expertise to obtain the intended outcomes, through interaction with the logic toolkit. In return, the RCC–5 alignments deliver a level of explicitness and verbal precision that exceeds that of any published alternative [4, 5, 6, 9, 16, 17]. That is, when we assert – and the reasoning validates as consistent - relationships such as 2015.Neoaves == 2014. Neoaves, 2015.Pelecaniformes < 2014.Pelecaniformes, or 2015.Aequorlithornites_Clade1 >< 2014.Passerea_Clade3, we can thereby not just verbalize these instances of congruence and conflict, but transparently document and therefore understand their provenance in a global concept relationship graph (Figs. 11 and 13). In other words, the RCC–5 alignments provide a logically tractable means to identify and also *explain* the extent of conflict.

From these identification and explanation functions we can derive novel data management and knowledge retrieval services. Example queries include the following. (1) Show all congruent regions of the alignment and their complete sets of clade concept labels. (2) Modify this query to only apply to alignment regions subsumed under one particular concept and source, such as 2014. Columbea. (3) For any subset region of the global alignment (e.g., 2015./2014.Australaves), show the lowest-level pairs of children that are sampled congruently, versus those that are sampled incongruently. (4) Highlight within such an alignment region all clade concepts for which parent coverage is relaxed, and which show congruence as a result of this action. (5) Highlight sets of concepts where incongruence is due to differential granularity (sampling), versus actual overlap. (6) Identify and rank concepts that participate in the greatest number of overlapping relationships (Table 3). (7) Identify and rank the longest chains of nested, overlapping concept sets (Fig. 12). (8) Highlight the congruent lowest-level concept pairs whose incongruent placement into higher-level regions causes the chains of overlap. (9) To further dissect the instances of overlap, list all split-concept resolution labels in complementary triplets {A*B, A\b, B\a}, and provide for each the two immediate children and (again) the set of lower-level, whole-concept resolution regions that are differentially distributed by the split (Fig. 13 and Table 4). (10) Identify clade names that are unreliable across the source phylogenies; viz. identical clade name pairs that participate in concept labels with an incongruent relationship, or different clade names whose concept labels have a congruent relationship (Table 5).

All of the above queries, and many others we could propose, variously depend on our specific RCC–5 representation and reasoning conventions (see preceding Section). We suggest here that they present a new foundation for building logic-based, machine-scalable data integration services for the age of phylogenomics. Conceptualizing node identity and congruence this way addresses a gap in current systematic theory that is not adequately filled by other syntactic solutions.

We have shown elsewhere that Linnaean homonymy and synonymy relationships are unreliable indicators of congruence when constraints such as parent coverage are involved in establishing a concept’s referential extension [14, 26, 32]. Indeed, Code-enforced Linnaean naming is purposefully and productively designed to fixate the meaning of names by ostension, while allowing the intensional components to remain ambiguous [21, 54, 55, 56, 57]. Yet this trade-off effectively shifts the burden of disambiguating varying intensionalities associated with Linnaean names onto an additional, interpreting agent – typically human experts. Our RCC–5 alignment approach can be viewed as no more than a way to formalize the disambiguation effort so that it attains machine-interpretability.

Similarly, the use of a particular kind or sets of *phyloreferences* [58, 59, 60] is unlikely to reconstruct an alignment such as that of 2015./2014.Pelecanimorphae (Fig. 7), which would require: (1) an elaborate notion of phyloreference *homonymy* and *synonymy* (e.g., 2015. Pelecanifores versus 2014.Pelecaniformes,); (2) node-based definitions with comprehensive inclusion/exclusion constraints (covering all terminals); and (3) synapomorphy-based definitions at higher levels to model the relaxation of coverage constraints when lower-level concepts are wholly exclusive of each other (see also Figs. 1–4). All of these functions may be feasible in principle with phyloreferences, provided (again) that human expertise is permitted to enact them. However, we believe it is fair to presume that phyloreferences were not designed in the main for bringing out granular extensional differences between node concepts across multiple phylogenies. As such, their services may be best utilized in cases where continuous concept evolution and persistent phylogenomic conflict are not expected to be so frequent as to become the main driver of an identifier/relationship design.

### The role of trained judgment

The two largest alignments of 2015./2014.Neornithes (without) / 2015./2014.Telluraves jointly entail 895 concepts and 95 instances of relaxed parent coverage, with corresponding articulations of parent node congruence (e.g., Figs. 3 and 4). These input constraints provide us with 97 congruent regions in the global alignment, of which 85 regions – all parent level, and propagating to the congruent root region 2015.Neornithes == 2014.Neornithes – are gained only because of our indirect modeling of intensional node definitions (see S10 and S11 File sets). This outcome is consistent with intuitive or algorithmically supported signals expressed (e.g.) in [4, 6, 11], while being logically consistent and verbally explicit about how these definitions contribute to node identity, congruence, and conflict.

The strong contingency of the alignment outcome on expert intentions is neither surprising nor trivial. We wish to explore this dependency more deeply here. For instance, Redelings and Holder [23: pp. 5-6] in regards to the OToL supertree assembly and conflict identification method: “Any approach to supertree construction must deal with the need to adjudicate between conflicting input trees. We choose to deal with conflict by ranking the input trees, and preferring to include edges from higher-ranked trees. The merits of using tree ranking are questionable because the system does not mediate conflicts based on the relative amount of evidence for each alternative. […] In order to produce a comprehensive supertree, we also require a rooted taxonomy tree in addition to the ranked list of rooted input trees. Unlike other input trees, the taxonomy tree is required to contain all taxa, and thus has the maximal leaf set. We make the taxonomy tree the lowest ranked tree. […] Our method must resolve conflicts in order to construct a single supertree. However, the rank information used to resolve conflicts is an input to the method, not an output from the method. We thus perform curation-based conflict resolution, not inference-based conflict resolution.”

We can gather immediately from this passage that the method is deeply dependent on expert input regarding the relative ranking of each input phylogeny and also that of the similarly generated OToL taxonomy [24]. We can furthermore demonstrate that these ranking choices can lead to inconsistent outcomes whenever the sequentiality of differentially sampled input trees determines how concordance and conflict are negotiated by the algorithms (Figs. 14 and 15). For instance, if the less densely sampled tree is prioritized in order, and the taxonomy is not suitable to accommodate all non-matching components of a succeeding, more densely sampled tree, then the former tree is likely to show higher conflict with the latter in comparison to an inverse input sequence. Not surprisingly, a(ny) globally applied judgment of priority among differentially sampled trees is a poor proxy for actually modeling individual node concept intensionality; yet the latter is necessary to make reliable, local decisions between (1) conflict due to differential granularity versus (2) conflict due to overlap.

We can now return to the challenge posed in the Introduction. How to build a synthetic, phylogenomic data environment that offers reliable node identities and relationships in the face of continuous advancement and also persistent conflict? Our answer, exemplified with the neoavian explosion use case, is novel in the following sense. Given that such a service is desirable, we show that achieving it fundamentally depends on making and expressing upfront empirical commitments about the intensionalities of clade concepts whose children are incongruently sampled. Without embedding these trained judgments into the alignment input, we would lose the 85 congruent parent regions that are recoverable only under relaxed parent coverage. We would furthermore lose the ability to distinguish the former from more than 340 alignment regions that are *not* congruent, and lose the power to express the nature of this residual conflict – granularity versus overlaps – and how to resolve it.

In other words, if indeed we intended to build a synthetic data environment in which the two author teams [1, 2] and others researching avian phylogenomics can logically present and integrate their diverging node concepts and related data, the first step will be to recognize that reliable conceptualizations of node identity within such a system just cannot be provided through some mechanical, ‘objective’ criterion. Instead, we need to make deliberate room within our design framework for an *inclusive* standard of objectivity that embraces trained judgment as an integral part of node concept intensionality [30]. In that sense, phylogenomic syntheses *are* inference-based (*contra* [23]), as well as purpose-driven, because our goal as integrative biologists – expressed through RCC–5 alignments – is to maximize intensional node congruence. There may not be a more reliable criterion for achieving this than expert-provided input constraints, which draw on diverse and complex theoretical knowledge of node and character identity in particular topological contexts [40, 43, 61]. Logic representation and reasoning can help render these constraints explicit and consistent, while exposing additional implicit constraints in the MIR. But logic cannot substitute the expert aligners' intensional aims and definitions.

The upshot of this outcome is that the logic of phylogenomic data integration forces us, in an important sense, to become experts about externally generated results that *conflict* with those which we may (currently) publish or endorse. It forces us to become experts of another author team’s node concepts, to the point where we are comfortable with expressing counter-factual statements regarding their intensionalities in light of incongruent child sampling. We suspect that this will require a rather profound but necessary adjustment in achieving a culture of synthesis in systematics that is no longer manages conflict in this suppressive way: “If we do not agree, then it is either our view over yours, or we just collapse all conflicting node concepts into polytomies”. In contrast, using RCC–5 alignments requires us to build up the following perspective: “We may or may not agree with you (likely we do not), but in either case we understand your phylogenomic inference well enough to express our dis-/agreements in a logic-compatible syntax, and are prepared to assert and refine articulations from our concepts to yours for the purpose of maximizing intensional node congruence”. Only then can we expect to also maximize the empirical translatability of biological data linked to diverging phylogenomic hypotheses.

We suggest here that shifting towards this latter attitude will be more challenging for the systematic community that providing the operational logic to enable scalable alignments.

Automation of certain workflow components is certainly possible, but is ultimately not the hardest bottleneck to overcome to attain scalability for this approach. Therefore, future designs for data environments capable of verbalizing phylogenomic conflict and synthesis need to reflect deeply on strategies that would incentivize a shift towards a culture where experts routinely and explicitly assess the intensionalities of node concepts of our peers. Stated more plainly, if we wish to track progress and conflict across our phylogenomic inferences, we first need to design a value system that better enables and motivates experts to do so.

## Author summary

Synthetic platforms for phylogenomic knowledge tend to manage conflict between different evolutionary reconstructions in the following way: “If we do not agree, then it is either our view over yours, or we just collapse all conflicting node concepts into polytomies”. We argue that this is not an equitable way to realize synthesis in this important biological domain. For instance, it would not be an adequate solution for building a unified data environment where multiple active author teams can endorse and yet also reconcile their diverging perspectives, side by side. Hence, we develop a novel system for *verbalizing* – i.e., consistently identifying and aligning – incongruent node concepts that reflects a more forward-looking attitude: “We may not agree with you, but nevertheless we *understand* your phylogenomic inference well enough to *express* our disagreements in a logic-compatible syntax, for the purpose of maximizing the empirical translatability of biological data linked to our diverging phylogenomic hypotheses”. We demonstrate that achieving such a notion of phylogenomic synthesis fundamentally depends on the application of trained expert judgment to stipulate parent node congruence in spite of incongruently sampled children. We have thereby outlined the core conditions for a more powerful language for integrating the evolving products of phylogenomic research.

## Author contributions

**Conceptualization**: Nico M. Franz, Bertram Ludäscher.

**Methodology**: Nico M. Franz, Shizhuo Yu, Bertram Ludäscher.

**Software**: Shizhuo Yu, Bertram Ludäscher, Joseph W. Brown.

**Validation**: Nico M. Franz, Lukas J. Musher, Joseph W. Brown.

**Visualization**: Nico M. Franz, Shizhuo Yu, Bertram Ludäscher, Joseph W. Brown.

**Writing – original draft**: Nico M. Franz.

**Writing – review & editing**: Nico M. Franz, Lukas J. Musher, Joseph W. Brown, Bertram Ludäscher.

## Acknowledgments

The authors thank Shawn Bowers, Andrew Johnston, Parisa Kianmajd, Jonathan Rees, Beckett Sterner, Guanyang Zhang for technical and conceptual input that improved the contents of this manuscript. Support of the authors/ research through the National Science Foundation is kindly acknowledged (NMF: DEB-1155984, DBI-1342595; JWB: DEB-1207915; BL: IIS-1118088, DBI-1147273).

## Supporting information captions

**S1A Text. Reasoner input constraints for the 2015./2014.Psittaciformes alignment, with coverage globally applied**. Includes information on run commands; and 0 instances of “no coverage”. File format: .txt.

**S1B Fig. Input visualization for the 2015./2014.Psittaciformes alignment, with coverage globally applied.** File format: pdf.

**S2A Fig. Alignment visualization for the 2015./2014.Psittaciformes alignment, with coverage globally applied**. File format: pdf.

**S2B File. Set of Maximally Informative Relations (MIR) inferred for the 2015./2014.Psittaciformes alignment, with coverage globally applied.** Total = 108 MIR. File format: .csv.

**S3A Text. Reasoner input constraints for the 2015./2014.Psittaciformes alignment, with coverage locally relaxed**. Includes information on run commands; and 4 instances of “no coverage”. File format: .txt.

**S3B Fig. Input visualization for the 2015./2014.Psittaciformes alignment, with coverage locally relaxed.** File format: .pdf.

**S4A Fig. Alignment visualization for the 2015./2014.Psittaciformes alignment, with coverage locally relaxed.** File format: .pdf.

**S4B File. Set of Maximally Informative Relations (MIR) inferred for the 2015./2014.Psittaciformes alignment, with coverage locally relaxed**. Total = 160 MIR. File format: .csv.

**S5A Text. Reasoner input constraints for the alignment of passeriform clade concepts (“Phylo2015”) sec. 2015.PEA with the corresponding classification concepts (“Class2015”) sec. Gill & Donsker (2015); including the (paraphyletic) Class2015.Eurylaimidae.** Includes information on run commands; and 0 instances of “no coverage”. File format: .txt.

**S5B Fig. Input visualization for the alignment of passeriform clade concepts (“Phylo2015”) sec. 2015.PEA with the corresponding classification concepts (“Class2015”) sec. Gill & Donsker (2015); including the (paraphyletic) Class2015.Eurylaimidae.** File format: pdf.

**S6A Fig. Alignment visualization for the alignment of passeriform clade concepts (“Phylo2015”) sec. 2015.PEA with the corresponding classification concepts (“Class2015”) sec. Gill & Donsker (2015); including the (paraphyletic) Class2015.Eurylaimidae**. File format: .pdf.

**S6B File. Set of Maximally Informative Relations (MIR) inferred for the alignment of passeriform clade concepts (“Phylo2015”) sec. 2015.PEA with the corresponding classification concepts (“Class2015”) sec. Gill & Donsker (2015); including the (paraphyletic) Class2015.Eurylaimidae**. Total = 63 MIR. File format: .csv.

**S7A Text. Reasoner input constraints for the alignment of tyrannoid clade concepts (“Phylo2015”) sec. 2015.PEA with the corresponding classification concepts (“Class2015”) sec. Gill & Donsker (2015); including the (paraphyletic) Class2015.Tityridae**. Includes information on run commands; and 0 instances of “no coverage”. File format: .txt.

**S7B Fig. Input visualization for the alignment of tyrannoid clade concepts (“Phylo2015”) sec. 2015.PEA with the corresponding classification concepts (“Class2015”) sec. Gill & Donsker (2015); including the (paraphyletic) Class2015.Tityridae.** File format: pdf.

**S7C Fig. Alignment visualization for the alignment of tyrannoid clade concepts (“Phylo2015”) sec. 2015.PEA with the corresponding classification concepts (“Class2015”) sec. Gill & Donsker (2015); including the (paraphyletic) Class2015.Tityridae**. File format: .pdf.

**S7D File. Set of Maximally Informative Relations (MIR) inferred for the alignment of tyrannoid clade concepts (“Phylo2015”) sec. 2015.PEA with the corresponding classification concepts (“Class2015”) sec. Gill & Donsker (2015); including the (paraphyletic) Class2015.Tityridae**. Total = 140 MIR. File format: .csv.

**S8A Text. Reasoner input constraints for the alignment of procellariiform clade concepts (“Phylo2015”) sec. 2015.PEA with the corresponding classification concepts (“Class2015”) sec. Gill & Donsker (2015); including the (paraphyletic) Class2015.Hydrobatidae and Class2015.Procellariidae**. Includes information on run commands; and 0 instances of “no coverage”. File format: .txt.

**S8B Fig. Input visualization for the alignment of procellariiform clade concepts (“Phylo2015”) sec. 2015.PEA with the corresponding classification concepts (“Class2015”) sec. Gill & Donsker (2015**); **including the (paraphyletic) Class2015.Hydrobatidae and Class2015.Procellariidae**. File format: pdf.

**S8C Fig. Alignment visualization for the alignment of procellariiform clade concepts (“Phylo2015”) sec. 2015.PEA with the corresponding classification concepts (“Class2015”) sec. Gill & Donsker (2015); including the (paraphyletic) Class2015.Hydrobatidae and Class2015.Procellariidae.** File format: pdf.

**S8D File. Set of Maximally Informative Relations (MIR) inferred for the alignment of procellariiform clade concepts (“Phylo2015”) sec. 2015.PEA with the corresponding classification concepts (“Class2015”) sec. Gill & Donsker (2015); including the (paraphyletic) Class2015.Hydrobatidae and Class2015.Procellariidae.** Total = 221 MIR. File format: .csv.

**S9A Text. Reasoner input constraints for the alignment of caprimulgiform clade concepts (“Phylo2015”) sec. 2015.PEA with the corresponding classification concepts (“Class2015”) sec. Gill & Donsker (2015); including the (paraphyletic) Class2015.Caprimulgiformes**.

Includes information on run commands; and 0 instances of “no coverage”. File format: .txt.

**S9B Fig. Input visualization for the alignment of caprimulgiform clade concepts (“Phylo2015”) sec. 2015.PEA with the corresponding classification concepts (“Class2015”) sec. Gill & Donsker (2015); including the (paraphyletic) Class2015.Caprimulgiformes.** File format: .pdf.

**S9C Fig. Alignment visualization for the alignment of caprimulgiform clade concepts (“Phylo2015”) sec. 2015.PEA with the corresponding classification concepts (“Class2015”) sec. Gill & Donsker (2015); including the (paraphyletic) Class2015.Caprimulgiformes**. File format: .pdf.

**S9D File. Set of Maximally Informative Relations (MIR) inferred for the alignment of caprimulgiform clade concepts (“Phylo2015”) sec. 2015.PEA with the corresponding classification concepts (“Class2015”) sec. Gill & Donsker (2015); including the (paraphyletic) Class2015.Caprimulgiformes**. Total = 672 MIR. File format: csv.

**S10A Text. Reasoner input constraints for the 2015./2014.Neornithes alignment (excepting 2015./2014.Telluraves), with coverage locally relaxed**. Includes information on run commands; and 58 instances of “no coverage”. File format: .txt.

**S10B Fig. Input visualization for the 2015./2014.Neornithes alignment (excepting 2015./2014.Telluraves), with coverage locally relaxed**. File format: pdf.

**S10C Fig. Alignment visualization for the 2015./2014.Neornithes alignment (excepting 2015./2014.Telluraves), with coverage locally relaxed**. File format: pdf.

**S10D File. Set of Maximally Informative Relations (MIR) inferred for the 2014./2014.Neornithes alignment (excepting 2015./2014.Telluraves), with coverage locally relaxed.** Total = 68,208 MIR. File format: .csv.

**S11A Text. Reasoner input constraints for the 2015./2014.Telluraves alignment, with coverage locally relaxed**. Includes information on run commands; and 37 instances of “no coverage”. File format: .txt.

**S11B Fig. Input visualization for the 2015./2014.Telluraves alignment, with coverage locally relaxed.** File format: .pdf.

**S11C Fig. Alignment visualization for the 2015./2014.Telluraves alignment, with coverage locally relaxed**. File format: .pdf.

**S11D File. Set of Maximally Informative Relations (MIR) inferred for the 2015. /2014.Telluraves alignment, with coverage locally relaxed**. Total = 32,864 MIR. File format: .csv.

S12A Text. Reasoner input constraints for the 2015./2014.Pelecanimorphae alignment, with coverage locally relaxed. Includes information on run commands; and 2 instances of “no coverage”. File format: .txt.

**S12B Fig. Input visualization for the 2015./2014.Pelecanimorphae alignment, with coverage locally relaxed**. File format: .pdf.

**S12C Fig. Alignment visualization for the 2015./2014.Pelecanimorphae alignment, with coverage locally relaxed**. File format: pdf.

**S12D File. Set of Maximally Informative Relations (MIR) inferred for the 2015. /2014.Pelecanimorphae alignment, with coverage locally relaxed.** Total = 200 MIR. File format: .csv.

**S13A Text. Reasoner input constraints for the 2015.Passeri/2014.Passeriformes_Clade2 alignment, with coverage locally relaxed. Includes information on run commands; and 1 instance of “no coverage”.** File format: .txt.

**S13B Fig. Input visualization for the 2015.Passeri/2014.Passeriformes_Clade2 alignment, with coverage locally relaxed.** File format: pdf.

**S13C Fig. Alignment visualization for the 2015.Passeri/2014.Passeriformes_Clade2 alignment, with coverage locally relaxed.** File format: pdf.

**S13D File. Set of Maximally Informative Relations (MIR) inferred for the 2015. Passeri/2014.Passeriformes_Clade2 alignment, with coverage locally relaxed. Total = 140 MIR.** File format: .csv.

**S14A Text. Reasoner input constraints for the 2015./2014.Telluraves alignment (higher-level subset), under whole-concept resolution. Includes information on run commands; and 0 instances of “no coverage”.** File format: .txt.

**S14B Fig. Input visualization for the 2015./2014.Telluraves alignment (higher-level subset), under whole-concept resolution.** File format: pdf.

**S14C Fig. Alignment visualization for the 2015./2014.Telluraves alignment (higher-level subset), under whole-concept resolution.** File format: pdf.

**S14D File. Set of Maximally Informative Relations (MIR) inferred for the 2015./2014.Telluraves alignment (higher-level subset), under whole-concept resolution.** Total = 81 MIR. File format: .csv.

**S15A Text. Reasoner input constraints for the 2015./2014.Telluraves alignment (higher-level subset), under split-concept resolution. Includes information on run commands; and 0 instances of “no coverage”.** File format: .txt.

**S15B Fig. Input visualization for the 2015./2014.Telluraves alignment (higher-level subset), under split-concept resolution.** File format: pdf.

**S15C Fig. Alignment visualization for the 2015./2014.Telluraves alignment (higher-level subset), under split-concept resolution.** File format: pdf.

**S15D File. Set of Maximally Informative Relations (MIR) inferred for the 2015./2014.Telluraves alignment (higher-level subset), under split-concept resolution. Total = 81 MIR.** File format: .csv.

**S16A Text. Reasoner input constraints for the 2015./2014.Neornithes alignment, under whole-concept resolution, ranging from the root to the ordinal level (with exceptions where needed). Includes information on run commands; and 4 instances of “no coverage”.** File format: .txt.

**S16B Fig. Input visualization for the 2015./2014.Neornithes alignment, under whole-concept resolution, ranging from the root to the ordinal level (with exceptions where needed).** File format: .pdf.

**S16C Fig. Alignment visualization for the 2015./2014.Neornithes alignment, under whole-concept resolution, ranging from the root to the ordinal level (with exceptions where needed).** File format: .pdf.

**S16D File. Set of Maximally Informative Relations (MIR) inferred for the 2015./2014.Neornithes alignment, under whole-concept resolution, ranging from the root to the ordinal level (with exceptions where needed). Total = 8,051 MIR.** File format: csv.

**S17A Text. Reasoner input constraints for the 2015./2014.Neoaves alignment, under whole-concept resolution, limited to the main conflict region. Includes information on run commands; and 0 instances of “no coverage”.** File format: .txt.

**S17B Fig. Input visualization for the 2015./2014.Neoaves alignment, under whole-concept resolution, limited to the main conflict region.** File format: pdf.

**S17C Fig. Alignment visualization for the 2015./2014.Neoaves alignment, under whole-concept resolution, limited to the main conflict region.** File format: pdf.

**S17D File. Set of Maximally Informative Relations (MIR) inferred for the 2015./2014.Neoaves alignment, under whole-concept resolution, limited to the main conflict region. Total = 441 MIR.** File format: .csv.

**S18A Text. Reasoner input constraints for the 2015./2014.Neoaves alignment, under split-concept resolution, limited to the main conflict region. Includes information on run commands; and 0 instances of “no coverage”.** File format: .txt.

**S18B Fig. Input visualization for the 2015./2014.Neoaves alignment, under split-concept resolution, limited to the main conflict region.** File format: pdf.

**S18C Fig. Alignment visualization for the 2015./2014.Neoaves alignment, under split-concept resolution, limited to the main conflict region.** File format: pdf.

**S18D File. Set of Maximally Informative Relations (MIR) inferred for the 2015. /2014.Neoaves alignment, under split-concept resolution, limited to the main conflict region. Total = 441 MIR.** File format: .csv.

**S19 File. Output tree file (visualized using FigTree) of the OToL conflict visualization method, with 2014.JEA as the primary source phylogeny and 2015.PEA as the alternative.** File format: .tre.

**S19 File. Output tree file (visualized using FigTree) of the OToL conflict visualization method, with 2015.PEA as the primary source phylogeny and 2014.JEA as the alternative.** File format: .tre.

